# An integrated homeostatic reinforcement learning theory of motivation explains the transition to cocaine addiction

**DOI:** 10.1101/029256

**Authors:** Mehdi Keramati, Audrey Durand, Paul Girardeau, Boris Gutkin, Serge Ahmed

**Author notes:** Correspondence should be addressed to M.K. or S.H.A.

## Abstract

Drugs of abuse implicate both reward learning and homeostatic regulation mechanisms of the brain. Theories of addiction, thus, have mostly depicted this phenomenon as pathology in either habit-based learning system or homeostatic mechanisms. Showing the limits of those accounts, we hypothesize that compulsive drug seeking arises from drugs hijacking a system that integrates homeostatic regulation mechanism with goal-directed action/behavior. Building upon a recently developed homeostatic reinforcement learning theory, we present a computational theory proposing that cocaine reinforces goal-directed drug-seeking due to its rapid homeostatic corrective effect, whereas its chronic use induces slow and long-lasting changes in homeostatic setpoint. Our theory accounts for key behavioral and neurobiological features of addiction, most notably, escalation of cocaine use, drug-primed craving and relapse, and individual differences underlying susceptibility to addiction. The theory also generates unique predictions about the mechanisms of cocaine-intake regulation and about cocaine-primed craving and relapse that are confirmed by new experiments.

**Significance:** Chronic use of addictive drugs renders increased motivation in planning to obtain and consume the drugs, despite their adverse social, occupational, and health consequences. It is as if addicts gradually develop a strong need for the drug and use their cognitive abilities and the knowledge of their environment in order to fulfil that need. In this paper, we build a mathematical model of this conception of addiction and show through quantitative simulations that such a model actually behaves in the same way that human addicts or laboratory animals that are exposed to cocaine behave. For example, the model shows gradually increasing motivation for drugs, relapse after long periods of abstinence, and individual differences in susceptibility to addiction.

## Introduction

Drug addiction or substance use disorder is a major public health problem. Many years of experimental research have revealed the complex, multi-faceted nature of this phenomenon. One dominant view in modern theories of addiction is that the pharmacological effects of the drugs, either through artificially over-reinforcing drug-seeking responses by hijacking the dopaminergic system (1–3), or through decreasing activity in prefrontal cortex (3), induces a transition from voluntary and goal-directed, to habit-based decision processes. Despite lack of direct evidence that compulsive drug seeking is a habitual behavior, this view has deeply shaped research on addiction. On the theoretical front, for example, previous computational models (2, 4–6) based on the Reinforcement Learning (RL) theory depicted addiction as maladaptive over-estimation of the (habitual) value associated with past drug-seeking behavioral responses. Repetitive over-reinforcement of such responses renders them insensitive to the possible adverse consequences associated with drugs and engendering compulsive drug seeking.

Challenging this perspective, we show that numerous key aspects of addiction cannot be explained by habits. In contrast, we demonstrate theoretically that a goal-directed system integrated with a homeostatic regulation (HR) mechanism provides a more complete and parsimonious account of core behavioral features of addiction, notably of those reproduced in animal models.

HR-based theories have in fact had a long, though less noticed history in theories of addiction. HR models (7, 8) axiomatize that animals’ objective is physiological stability. To this end, corrective responses are triggered when a deviation of some key physiological variables from their hypothetical setpoints is sensed. HR-based models of addiction assume that drugs modulate the level of some homeostatically-regulated internal variables. Drug-seeking response, thus, is elicited when drug can reduce the homeostatic deviation of those variables (9–11). This class of models successfully explains the regular pattern of drug self-administration (12, 13) (SA) in animal models of addiction, attributing it to the animal’s desire for defending homeostasis by regularly compensating for the depleted drug level in the brain. However, lack of associative learning systems fundamentally limits the explanatory power of those theories.

Addressing this general shortfall of HR models, and taking into account the apparent coupling of the brain reward learning and homeostatic regulation mechanisms (14), a recently proposed homeostatic reinforcement learning (HRL) theory (15) provided a mathematical framework for interaction between the homeostatic and learning systems. The HRL theory proposes that the rewarding value of an action stems from the approximated capacity of that action’s outcome to reduce the homeostatic deviation of the organism. This computed reward is then used by associative learning mechanisms as a source of reinforcing associations.

Built upon this framework, our integrative theory of addiction incorporates a minimal model of the pharmacological effect of cocaine on several components of the HRL (15) system. Crudely speaking, we argue that addiction stems from the drugs hijacking a goal-directed reward-learning process that aims at fulfilling the physiological needs of the organism.

Critically, we propose that the acute effect of cocaine alters the level of an internal variable that is regulated homeostatically under normal (drug-free) conditions. We further propose that chronic drug-use progressively alters the setpoint level of this variable, through drug-induced plasticity mechanisms. Simulating the model that incorporates these assumptions into the HRL theory, we replicate and explain potential mechanisms underlying a wide range of behavioral and neurobiological data from rat (13, 16–22), monkey (23, 24), and human (25) experiments on cocaine addiction. Furthermore, the model makes several new testable predictions that contrast with the prediction of other models. We present new experimental data that confirm several of these unique predictions, notably regarding the exact mechanisms by which cocaine intake is regulated. Future research will be required to test the merit of the other predictions. Our theory argues that addiction is a consequence of long-lasting drug-induced plasticity in the brain HRL system, thereby identifying potential targets for addiction therapy.

## RESULTS

### Theory sketch

The basis of our theory stems from the computational framework of Homeostatic Reinforcement Learning (15) (HRL). The HRL theory, like most neuroeconomic theories, postulates that animals learn, and then exploit the learned environmental contingencies in order to maximize attainment of rewarding outcomes. Uniquely to HRL, when an outcome affects a homeostatically regulated variable, its rewarding value is defined by its ability to fulfill the homeostatic needs of the organism. In other words, the homeostatic-based primary reward is measured by the outcome-induced anticipated reduction in the deviation of the internal physiological state from the homeostatic setpoint. Consequently, HRL model has been proven to provide sufficient machinery for defending homeostasis through reward maximization (15).

Our theory of addiction is based on the critical hypothesis that cocaine acts on an internal variable (hereafter denoted by *h_t_*) that is regulated by the HRL mechanism. We assume in the model that in proportion with striatal cocaine concentration, *h_t_* elevates initially upon infusion of the drug, and then falls gradually as cocaine is progressively eliminated from the site-of-action (Fig. 1B). Thus, when *h_t_* is below its setpoint level (denoted by *h*^*^), cocaine infusion results in the reduction of homeostatic deviation (unless it induces an overshoot) proportional to the self-administered dose, rendering cocaine outcome rewarding (Fig. 1C). This rewarding value becomes the source of reinforcement of cocaine-seeking behavior, which is supported by an action-outcome associative learning mechanism (i.e., goal-directed decision process) (Fig. 1D). We later postulate, and discuss supporting evidence, that this internal variable (*h_t_*) is a monotonically increasing function of the tonic striatal dopamine (DA) concentrations. That is, as cocaine directly increases striatal DA level (26), it elevates the internal state *h_t_*. We assume that this drug-controlled internal state, aside from its deviation-reduction effect, also act on the state-recognition system, analogous to the role of contextual states. Simply put, in parallel with external stimuli, cocaine level also influences the state of the world that the animal finds itself in. More precisely, the state-space in which the animal learns and exploits instrumental associations is an augmented space composed of both internal and external states (). This assumption is supported by many studies showing that cocaine, like other drugs of abuse, acts as an interoceptive and discriminative cue (27–29).

**Fig. 1.**
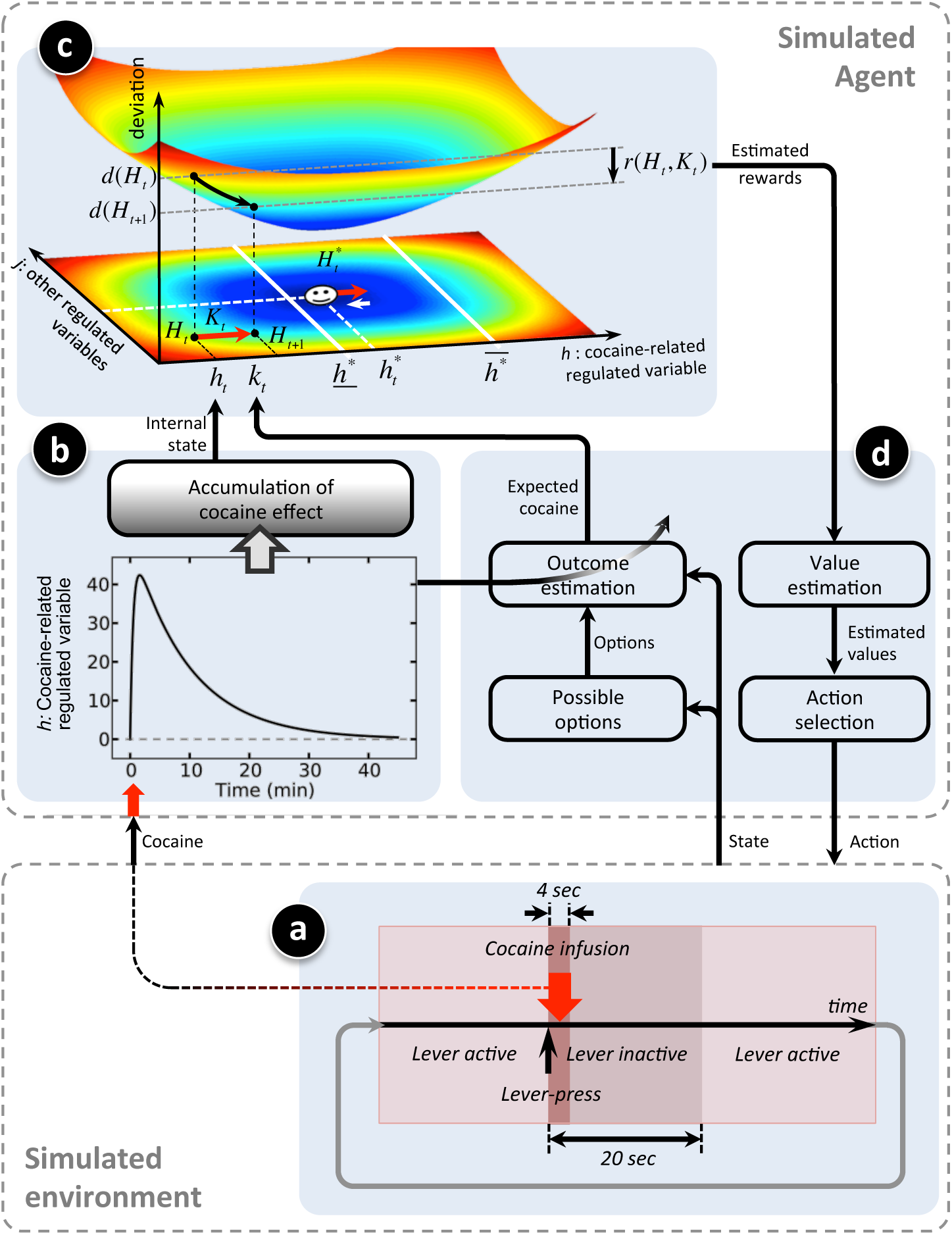
Schematics of the model. (A) Consisted with replicated experiments (9, 13, 16–19, 21, 22, 33, 34), the model is simulated in a self-administration paradigm where each lever-press (fixed-ratio one) initiates an intravenous infusion of cocaine over 4sec, followed by a 20sec time-out period during which the lever is inactive. (B) A certain brain internal variable (h) initially elevates upon a single infusion and then falls gradually as circulating cocaine degrades. The dynamics of *h* is compatible with cocaine-induced pharmacodynamics of tonic dopamine in the NAc (20). The effect of consecutive infusions accumulates over time. (C) The rewarding value (*r*) of an outcome (e.g. a certain dose of cocaine, indicated by *K_t_*) is equal to its ability in decreasing the distance (drive, indicated by *d*(*H_t_*)) of the internal state (*H_t_*) from the homeostatic setpoint (*H*^*^). *H_t_* is a vector composed of *h_t_* and other homeostatically-regulated variables. In parallel with this acute effect, every infusion also triggers a slow adaptive mechanism that slightly shifts the setpoint forward, capturing down-regulation of D2 receptors after chronic cocaine use. Absence of cocaine results in slow recovery of the setpoint to its initial level. 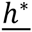 and 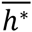 indicate the lower and upper bounds of the setpoint, respectively. (D) Given its current external state, the agent predicts the expected outcome of each possible choice and based on that, estimates the drive-reduction rewarding values of the choices. According to the estimated values, the agent selects an action. The curved arrow represents updating outcome expectancies based on feedbacks received from the environment.

In parallel with the acute effect of cocaine, chronic cocaine use induces several long-lasting neural plasticity, including the down-regulation of D2 receptors availability (30, 31) and reduced dopamine release in the striatum (32). These slow processes can be captured by an adaptive plasticity mechanism where the setpoint gradually shifts to compensate for the drug-induced excessive DA concentration (discuss later). In this respect, we simply assume that every infusion of cocaine results in a dose-dependent elevation of the setpoint. Also, abstinence gradually lowers back the setpoint to its initial level (Fig. 1C).

### Behavioral simulations

There is now strong evidence showing that following a history of extended access to cocaine SA, rats present behavioral changes that recapitulate important behavioral features of cocaine addiction. It was demonstrated in a seminal experiment (13, 16) that with one hour of access per session (short access or ShA) to intravenous cocaine SA, drug intake remained low and stable. In contrast, with 6 hours of access (long access or LgA), drug intake gradually escalated over days and eventually reached a level 200% greater than that of ShA rats.

Consistent with replicated experiments (9, 13, 16–19, 21, 22, 33, 34), we simulated the model in a virtual task () where each lever-press (fixed-ratio one) initiated an intravenous infusion of cocaine over 4sec, followed by a 20sec time-out period during which the lever was inactive (Fig. 1A). Simulation results (Fig. 2) replicated experimental data (13, 16) (,). As the simulated agents starts each session in a cocaine-deprived state (internal state = *h_t_* = 0), a burst of responding (known as “loading”) occurs at the beginning of sessions in order to reach the setpoint. Afterward, agents take a steady level of responding to maintain homeostasis. Note that, as in the replicated experiments, the structure of the task was learned during a pre-training period. Also note that in all the simulations in this paper, one single set of values for the free parameters of the model is used ().

**Fig. 2.**
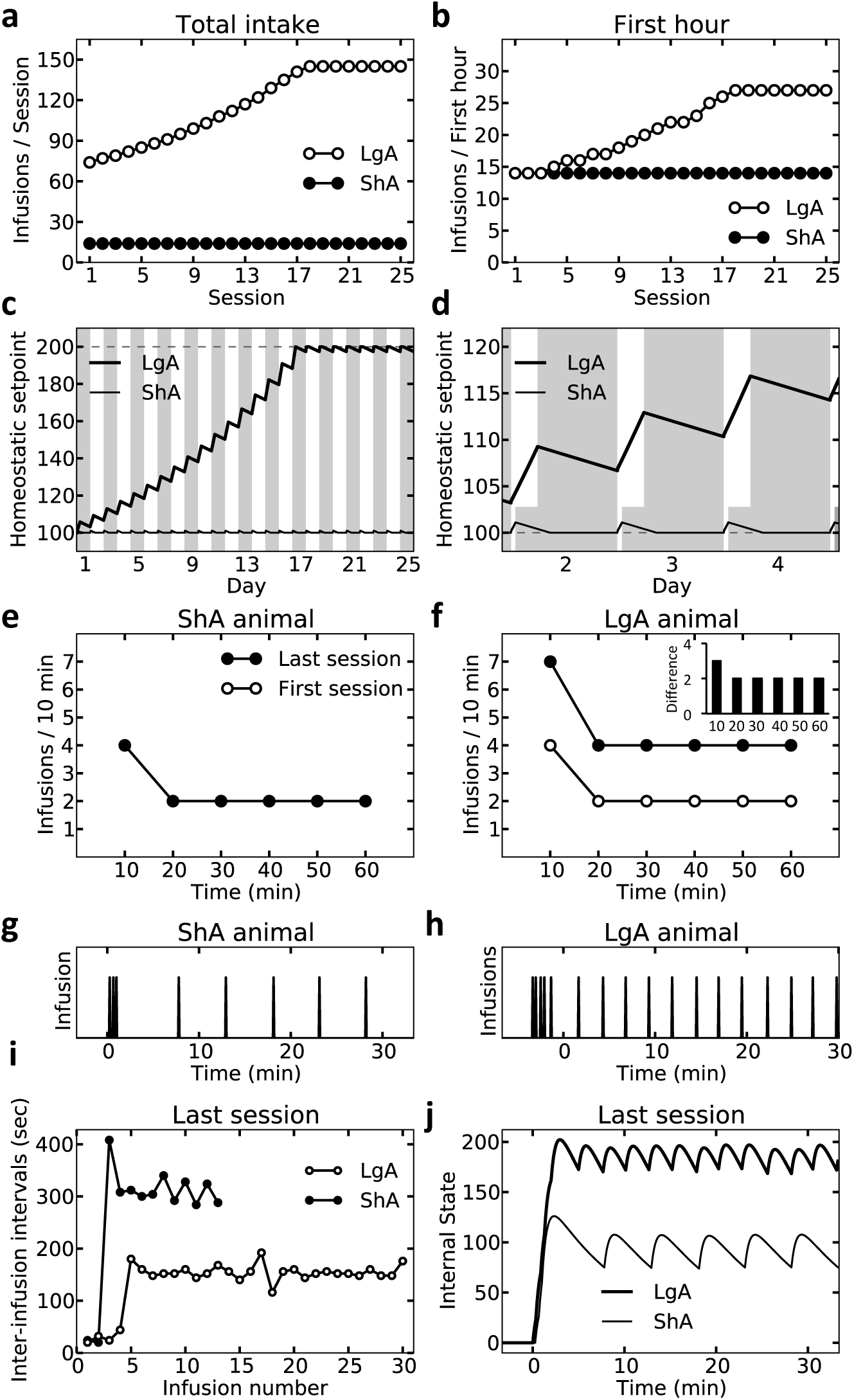
Simulation results replicating experimental data (13, 16) ( and ) on the escalation of cocaine self-administration under LgA condition. Starting from equal infusion rates in both groups, the LgA agent takes progressively more infusions than the ShA agent (A, B), due to gradual mounted level of the homeostatic setpoint (C). Focusing on the first few trials (D) shows that the elevation of the setpoint during a 6hr period cannot be recovered during the rest of the day (shaded area; 18hrs). Comparing the first and last sessions, the escalation pattern is also observable in the increased level of infusion during 10min bins in LgA (F), but not ShA (E) agents. In the LgA agent, this increase is stronger in the first, compared to other 10min bins (the inset in plot F; last minus first session). The rate of infusion in the first 10min block is greater than its steady level, in both ShA (E, G) and LgA (F, H) agents. This so-called “loading effect” is because the agents start the sessions in a cocaine-depleted internal state and thus, reaching the setpoint for the first time (J) requires several infusions with the least possible inter-infusion interval; i.e. 20 seconds (I).

Simulations fully replicate the pattern of escalation of infusion rate in LgA animals and make apparent that this escalation is due to the gradual elevation of the setpoint over several 6-hr access daily sessions. In ShA animals, however, setpoint elevation during every 1-hr session is small enough that the rest of the day (23-hrs) is sufficient for full recovery to the initial setpoint level. An elevated homeostatic setpoint in LgA agents induces escalation because maintaining the internal state at a higher setpoint requires more infusions to compensate for the relatively faster elimination of cocaine at higher concentrations. Clearly, at higher concentrations, a constant elimination rate (or half-life) naturally leads to higher total amount of eliminated cocaine.

Consistent with experimental data (), loading occurred in both ShA and LgA agents (Fig. 2E-H). Moreover, although infusion rate escalated in LgA agent at all 10-min bins, this effect was stronger at the first bin of sessions (the insets in F and D). Also, experimental data shows that the loading and maintenance phases are separated by a pause period during which no injection is taken (9). According to our model, the short delay between cocaine infusion and getting its full effect on the internal state (the ascending limb of the curve in Fig. 1A) causes the agents to keep taking cocaine for some extra times even after having taken enough for reaching the setpoint. This ensues overshooting the setpoint and thus, the pause phase occurs so that this excess cocaine degrades or washes out. In this respect, the model predicts that both loading and pause patterns will be more pronounced by decreasing the timeout period (). Intuitively, a shorter timeout provides the agent with the opportunity of adopting a higher infusion rate. This increases the number of extra infusions during the loading phase after the agent has already taken enough cocaine for reaching the setpoint. This results in a stronger overshoot and thus, a longer pause for offsetting it.

Experiments show that Post-escalation infusion rate, as well as the total amount of consumed cocaine per hour, are higher in LgA than ShA animals, for all unit doses of cocaine. However, whereas the infusion rate decreases as a function of dose, the amount consumed varies little (13, 16) (). Decreased response rate at higher doses is in contradiction with the classical RL theories, as they define utility (reward) functions that are increasing with respect to the outcome magnitude (35, 36). According to our model, however, the ultimate objective of the RL system is minimizing deviations from the setpoint. Therefore, increasing the dose induces lower infusion rate (Fig. 3A), but constant amount of consumption (Fig. 3B), so that the internal state fluctuates closely around the setpoint.

**Fig. 3.**
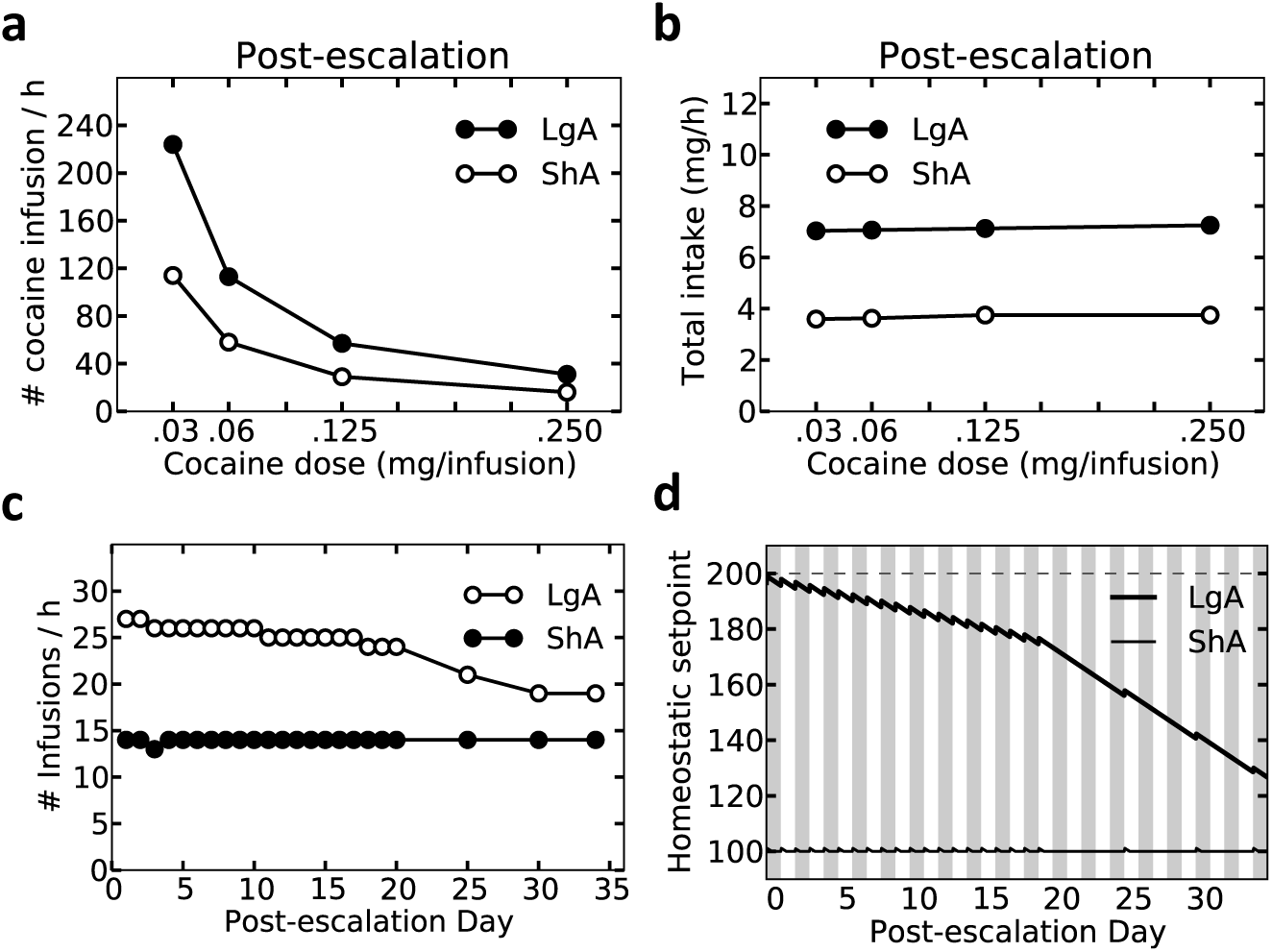
Simulation results replicating experimental data () on post-escalation dose effect on infusion rate and the total intake of cocaine (13, 16), as well as post-escalation reduced availability of cocaine (16). (A, B) Post-escalation infusion rate and the total amount of consumed cocaine per hour are higher in LgA than in ShA agents, for all unit doses of cocaine. However, whereas the infusion rate decreases as a function of dose (A), the amount consumed does not change with dose (B). (C, D) After escalation, both LgA and ShA agents are given limited (1hr/day) access to cocaine self-administration. This results in gradual recovery of the setpoint in the LgA agent (C) and thus, in decreasing the rate of infusion (A). After day 20, as in the experiment (16), the agents are given only 1hr access to cocaine in every five days. This speeds up the recovery process of the setpoint (B) and thus, accelerates the decreasing trend of infusion rate (A).

Evidence further shows that post-escalation reduction of access duration from 6hr to 1hr per day in LgA rats results in a gradual decline of the infusion rate. This decline becomes faster when access to cocaine is limited even more to only 1hr per week (16) (). According to our model, the shortened access allows the escalated setpoint to gradually return to its initial level (Fig. 3D), resulting in the infusion rate to reduce gradually (Fig. 3C). Although 1hr weekly access speeds up this process even further (Fig. 3C, D), five weeks is still insufficient for complete recovery to the initial, pre-escalation setpoint (Fig. 3D).

Given that access duration is a continuous parameter, one might expect that animals would show at least some small degree of escalation even at short access sessions. Yet strikingly, experiments revealed a minimum duration of access to the drug, below which no escalation is possible and above which the speed of escalation increases with the duration of access (17) (). Our model naturally accounts for critical duration; increasing the session duration prolongs the daily elevation and shortens the daily recovery periods of the setpoint level and thus, accelerates escalation (Fig. 4). For very short session durations (e.g. 1hr), however, the recovery effect during the rest of the day (e.g. 23hrs) completely cancels out the small elevation during the SA session. Above a certain critical duration, the two effects are no longer canceled, hence the escalation. The model predicts that at this critical session duration (3hr, in our simulations), the recovery period is just enough to cancel the cocaine-induced setpoint elevation (Fig. 4D). Hence, although such a session duration will not induce escalation (Fig. 4), it will preserve the infusion rate at an escalated level (e.g. if it has been escalated previously, using a 6hr access condition)().

**Fig. 4.**
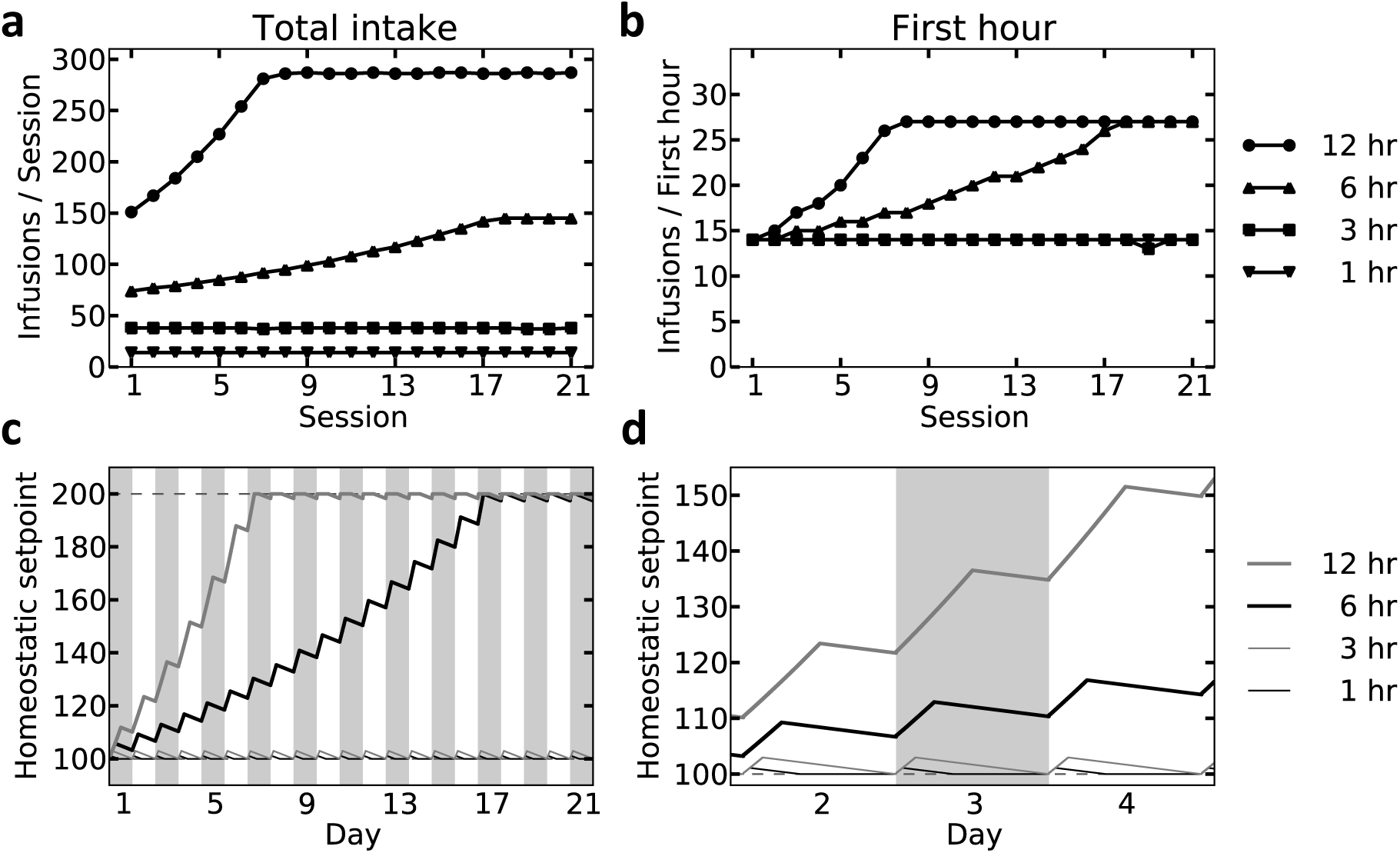
Simulation results replicating experimental data (17) () on the effect of session-duration on escalation. 1hr and 3hr daily access to cocaine self-administration do not induce escalation (A, B) since even under 3hr access, the elevation of the setpoint is cancelled out during the rest of the day (C, D). Rate of cocaine self-administration increased under 6hr and 12hr access conditions, and this increase was faster in the latter, than in the former (A, B).

Experimental results also show that post-escalation pairing of cocaine infusion with electric shock results in rapid suppression of responding within the first 45-minute session in both ShA and LgA rats (18). When the punishment is removed after this session, whereas LgA animals rapidly resume infusion with the pre-punishment rate within the first 45-min session, the infusion rate in ShA rats remains suppressed even after three sessions (18) (). This resistance to the long-term effects of punishment in LgA rats is supposed to capture compulsive drug seeking, which is the hallmark of addiction in humans. Our model reproduces these data (Fig. 5). According to the model, the high cost of lever-press during the electric-shock phase suppresses agents’ motivation for defending homeostasis. That is, the agents significantly reduce the response rate and press the lever only when the internal state falls far below the setpoint and thus, the setpoint deviation-reduction reward of cocaine outweighs the punishment (Fig. 5A). During post-punishment sessions, the homeostatic deviation in elevated-setpoint agents (LgA) is just high enough that motivates occasional lever-presses. Through this exploratory-like behavior the agents learn that the punishment is removed and therefore, resume SA with the pre-punishment rate. In ShA agents, however, the chance of exploring the punishment-removed environment is much less, due to relatively lower need for cocaine. Therefore, as with experimental results (18), a fraction of agents never try even a single lever-press, whereas some others try and thus learn that punishment is removed. As a result, and consistent with experimental data (18) (), ShA agents on average, but with a higher inter-individual variability, continue cocaine-seeking at a suppressed rate (Fig. 5B; see for details).

**Fig. 5.**
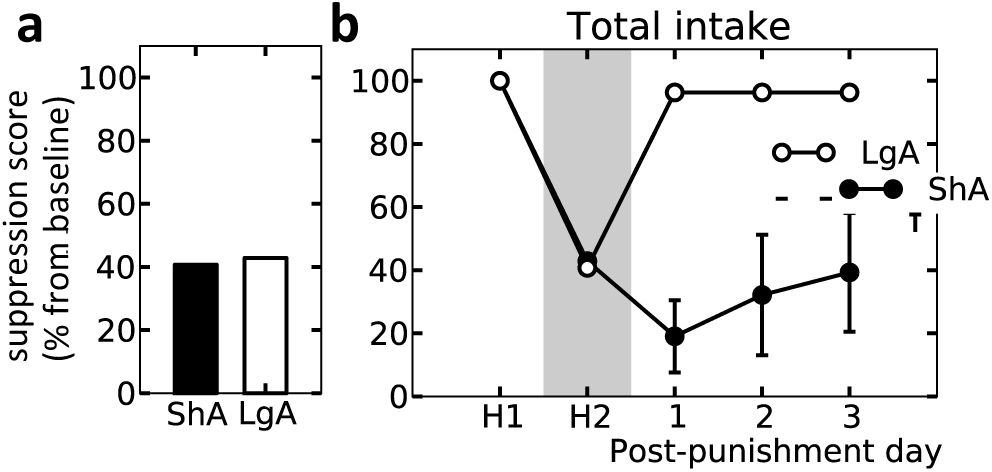
Simulation results replicating experimental data (18) () on the effect of extended drug-access on the punishment-induced suppression of cocaine seeking. After 25 days of 6hr vs. 1hr access to cocaine (Fig. 2), both LgA and ShA agents are provided with five sessions of 45min access to cocaine. Only in the second session (indicated by H2 in panel B) cocaine is paired with a punishment. This punishment results in equal rates of suppression (from baseline, indicated by H1) of cocaine self-administration in LgA and ShA agents (A). Whereas the LgA agent rapidly resumes self-administration after removal of the punishment, the ShA agent refrains during at least three consecutive days (B).

Perhaps the tell-tail hallmark of addiction is drug craving and relapse even after long-term abstinence. Well-validated animal models of craving and relapse show drug-primed reinstatement of cocaine seeking following extinction and abstinence. In one experiment (19), following 32 sessions of 6hr access to cocaine, rats underwent 10 days of reinstatement procedure. Each day consisted of 5 consecutive 45min blocks during which pressing the lever had no consequence (extinction). At the beginning of each block, rats received a single priming injection of cocaine with the following doses: 0, 0, 0.25, 0.5, and 1mg. Simulating the model (Fig. 6) in these same conditions reproduced many patterns observed in experimental results (19) ().

**Fig. 6.**
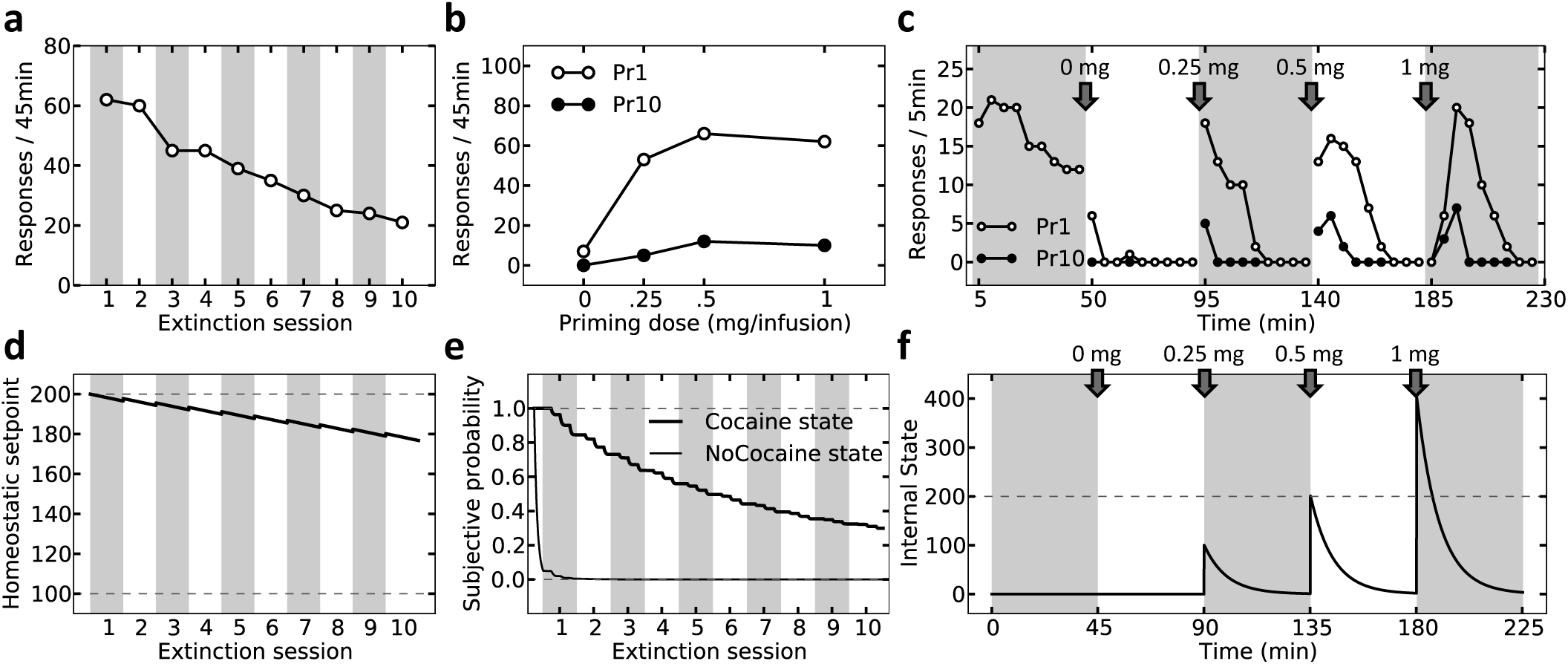
Simulation results replicating experimental data (19) () on the extinction of priming-induced reinstatement. After 25 days of 6hr access to cocaine (Fig. 2), the LgA agent undergoes a priming-induced reinstatement procedure during 10 consecutive days. Each day consists of 5 consecutive sessions of 45min during which pressing the lever has no consequence (extinction). At the beginning of each session, the agent receives a single priming injection of cocaine with the following doses: 0, 0, 0.25, 0.5, and 1mg. (A) The rate of lever-press in the 45min upon infusion of the highest dose (1mg) decreases progressively. (B) Such extinction also happens for other dose of cocaine, by comparing the first (Pr1) and the last (Pr10) extinction sessions. (C) Zooming on response rates at 5min intervals shows a more precise pattern of priming induced reinstatement. The escalated setpoint recovers only slightly during the 10 days of the experiment (D). 200 is the escalated level, and 100 is the initial normal level of the setpoint. Extinction of responding over 10 days is due to the decreased subjective probability of receiving cocaine, either when the agent is under cocaine, or when it is not (E). The extent to which the agent is under cocaine at any time-point (F) replicates experimental data (20) d and E). The dashed line indicates the setpoint level on the first day of the reinstatement experiment.

Firstly, as in behavioral data, although drug-seeking was extinguished in the first two blocks (no cocaine) of the first day, it relapsed dose-dependently and transiently after each priming injection, and was followed by a gradual return to pre-priming levels of cocaine seeking (Fig. 6C and C, curve Pr1). As mentioned before, the internal state in our model contributes to the mental representation of the agent’s state-space (). Thus, having been extinguished in a drug-free state during the first two blocks, the association between lever-pressing and cocaine has remained intact in the “under-cocaine” state (E, session 1). That is, the agent still expects cocaine in the “under-cocaine” state. Therefore, since priming injection re-induces the interoceptive stimuli of cocaine, drug-seeking relapses (B). However, as the priming cocaine degrades, the agent gradually returns to the “drug-free” state (F) where drug-seeking had been extinguished before. This explains why the priming effect on cocaine seeking is transient (C and C, curve Pr1).

Moreover, consistent with experimental results (19) (C), when the priming dose is very high (1mg), reinstatement does not occur instantaneously after injection, but peaks with a delay of 5–10min. According to our model, this is because upon injection of a high dose, the internal state overshoots the setpoint (Fig. 6F). This results in the agent entering the “under cocaine” state where the association between lever-press and outcome is still strong. However, the outcome of this state is not rewarding, as it will only increase the homeostatic deviation even further. Thus, the agent waits until the internal state sufficiently drops below the setpoint and then starts seeking cocaine (Fig. 6C).

Priming-induced reinstatement is in fact a transient phenomenon, and gradually extinguishes over approximately 10 days of experiencing the reinstatement procedure (19) (A, B). According to our model and consistent with previous ideas (37), the extinction of cocaine-induced reinstatement is due to gradual extinction of lever-cocaine associations in the “under-cocaine” state (Fig. 6E). This happens after the agent sufficiently experiences that association when it is under the effect of priming cocaine. Of critical clinical importance, our model predicts that the extinction of cocaine-induced reinstatement in this experimental procedure does not reflect complete recovery from addiction, as the setpoint is still elevated after 10 days of extinction (Fig. 6D). Thus, we predict that once rats are again given access to cocaine SA after the 10-day extinction procedure, the infusion rate must return rapidly (within only one 6hr session) to its escalated level (). This is because the setpoint remains at an elevated level, and the extinguished lever-cocaine associations can be re-learned within less than one hour (C), resulting in rapid re-escalation of infusion rate (A, B).

Critically, Long access to the drug not only causes escalation of dose, but also leads to a more pronounced cocaine-induced reinstatement of the drug-associated behaviors across all tested doses (21) (). According to our model, this is because the elevated setpoint level in the LgA agent induces a stronger homeostatic deprivation (deviation) and thus, results in a higher estimated value for the cocaine outcome. This, in turn, results in more pronounced reinstatement in the LgA agent ().

### Tested predictions

In our theory, the objective is to maintain the internal state as close as possible to the setpoint. To this end, the agent self-administers cocaine with a stable rate so that the internal state fluctuates regularly around the setpoint (Fig. 7A). Thus, a self-administration response is triggered each time the internal state drops sufficiently below the setpoint. This is in contrast to the previous regulatory models of cocaine addiction (9–11) where a response is triggered as soon as the internal state hits the setpoint (Fig. 7E). Therefore, those models predict that for different unit doses of cocaine, the response-triggering state will be the same (equal to the setpoint) (A, B), while our model predicts different triggering states for different unit doses (C, D). That is, the smaller the dose is, the closer to the setpoint the internal state will be maintained and thus, the higher the minimum level of cocaine concentration (or striatal DA level) will be.

**Fig. 7.**
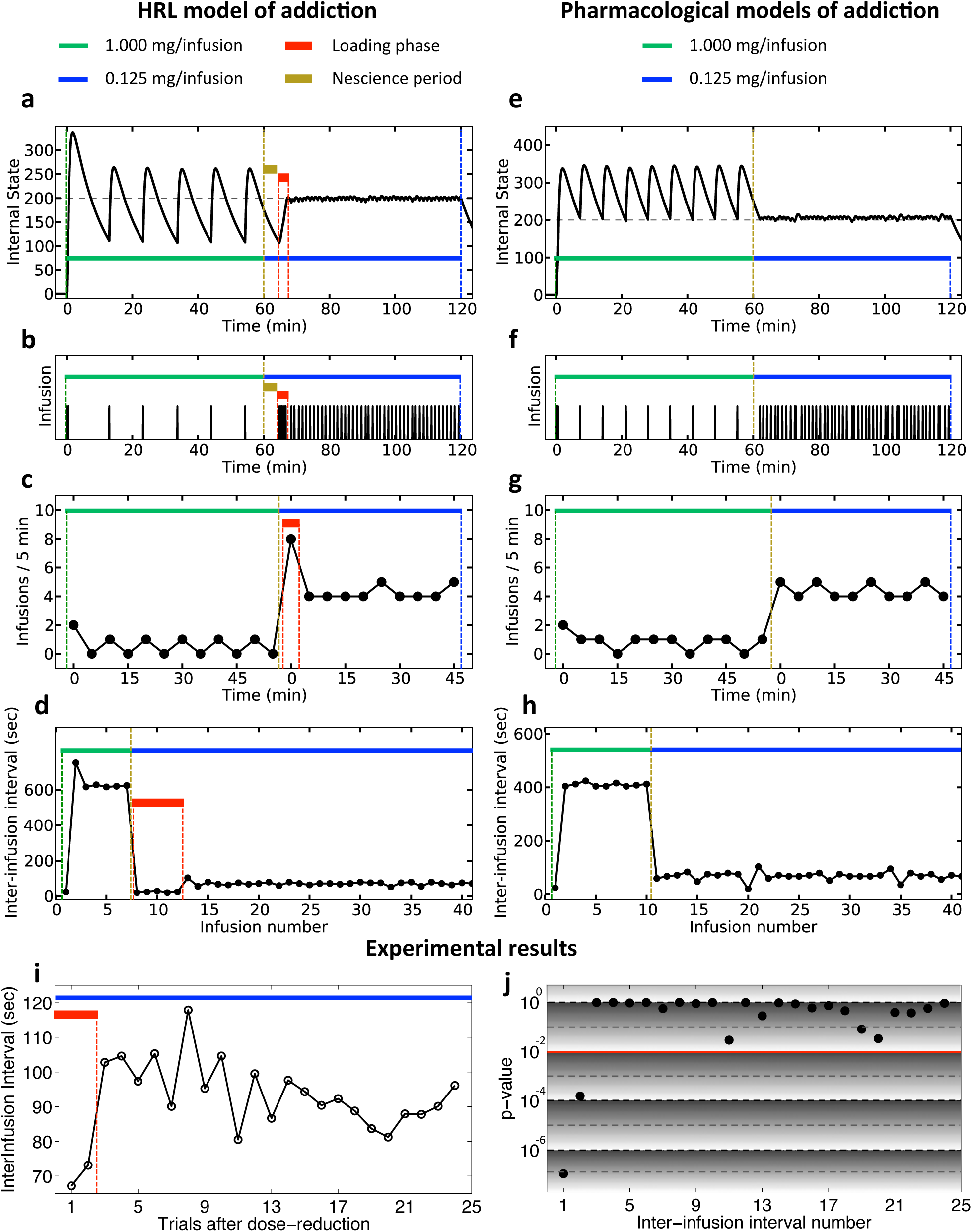
Simulation and experimental results on the effect of within-session reduction of unit-dose on self-administration pattern. After pre-training, both the simulated agents and the rats were tested in a two-hour session where each lever-press resulted in receiving a high vs. low dose of cocaine during the first vs. the second hour of the session. (A-D) Our model predicts a transitory burst of infusion rate after the dose reduction. In fact, when the dose is reduced, the agent still waits for a period (nescience) equal to previous inter-infusion intervals so as the internal state sufficiently drops below the setpoint. This is because the agent’s objective is oscillate around the setpoint in order to minimize deviations. Upon the first post-reduction response, the agent realizes the change and thus, responds intensively in order to catch the setpoint by shortened steps. After the setpoint is reached, the agent responds with a steady rate just to oscillate around the setpoint. (E-H) Previous models, in contrast, predict no response burst after dose reduction. In those models, a SA response is elicited every time the internal state drops below the setpoint. After dose-reduction, therefore, as soon as the internal state hits the setpoint, the agent starts responding with a new steady level. Confirming the prediction of our model, experimental results from rats (n=21) showed two significantly shorter inter-infusion intervals (III) right after the dose reduction, as compared to the later IIIs. (I) Average post-reduction III over all rats, over the latest three sessions. (J) *p*-values of one-sided t-tests with the alternative hypothesis that the *i*-th post-reduction III is less than the IIIs between the 10^th^ and the 20^th^ post-reduction infusions (when the response rate is supposedly converged to its new steady level).

While testing directly these diverging predictions () would require measurements of cocaine or DA levels immediately before each triggered response, they can nevertheless also be tested using a rather simple behavioral experiment involving a large, non-signaled within-session decrease in the unit dose of cocaine, from 1 to 0.0625 mg per injection. According to our model, the agent, initially unaware that the dose has been decreased, will wait until the internal state drops sufficiently below the setpoint. Then upon the first infusion of the lower dose, the agent will realize that the dose is reduced and that taking it is not enough to reach the setpoint. As a result, the agent will continue responding at the highest possible rate until the setpoint is reached (loading phase). This burst of responding will then be followed by a relatively lower response rate so as to fluctuate around the setpoint with the reduced dose (Fig. 7A-D). Previous models, in contrast, predict no loading phase after dose reduction, since the internal state is always maintained above the setpoint and thus, there is no undershooting to be compensated for (Fig. 7E-H).

These behavioral predictions have been tested in rats (n=21) trained in 2-hour sessions of cocaine SA with unit doses of 1 and 0.125 mg/injection during the first and the second hours, respectively. Results showed that the first two inter-infusion intervals (III) after reducing the dose were significantly shorter than IIIs between the 10^th^ and the 20^th^ post-reduction responses (Fig. 7I, J). This dose reduction-induced loading phase verifies our prediction and contradicts the prediction of previous models.

A further prediction of our model concerns individual differences in dose-response curves, arising from different setpoint levels (e.g. due to different levels of D2 receptors in the striatum, see discussion). Our theory predicts that animals are not motivated to respond for cocaine when its unit dose is less than a critical dose (dashed vertical line in Fig. 8A-F). This is because the costs associated with SA (cost of pressing the lever, etc.) outweigh the small drive-reduction effect of small doses of cocaine. However, this critical dose will be lower in animals with higher setpoint levels, since even a small dose has a great setpoint deviation-reduction effect for animals with a high setpoint (Fig. 8A-F). In fact, having a higher setpoint is equivalent to having a higher deprivation level, and deprivation level, in the HRL theory, has an excitatory effect on the setpoint deviation-reduction reward (15).

**Fig. 8.**
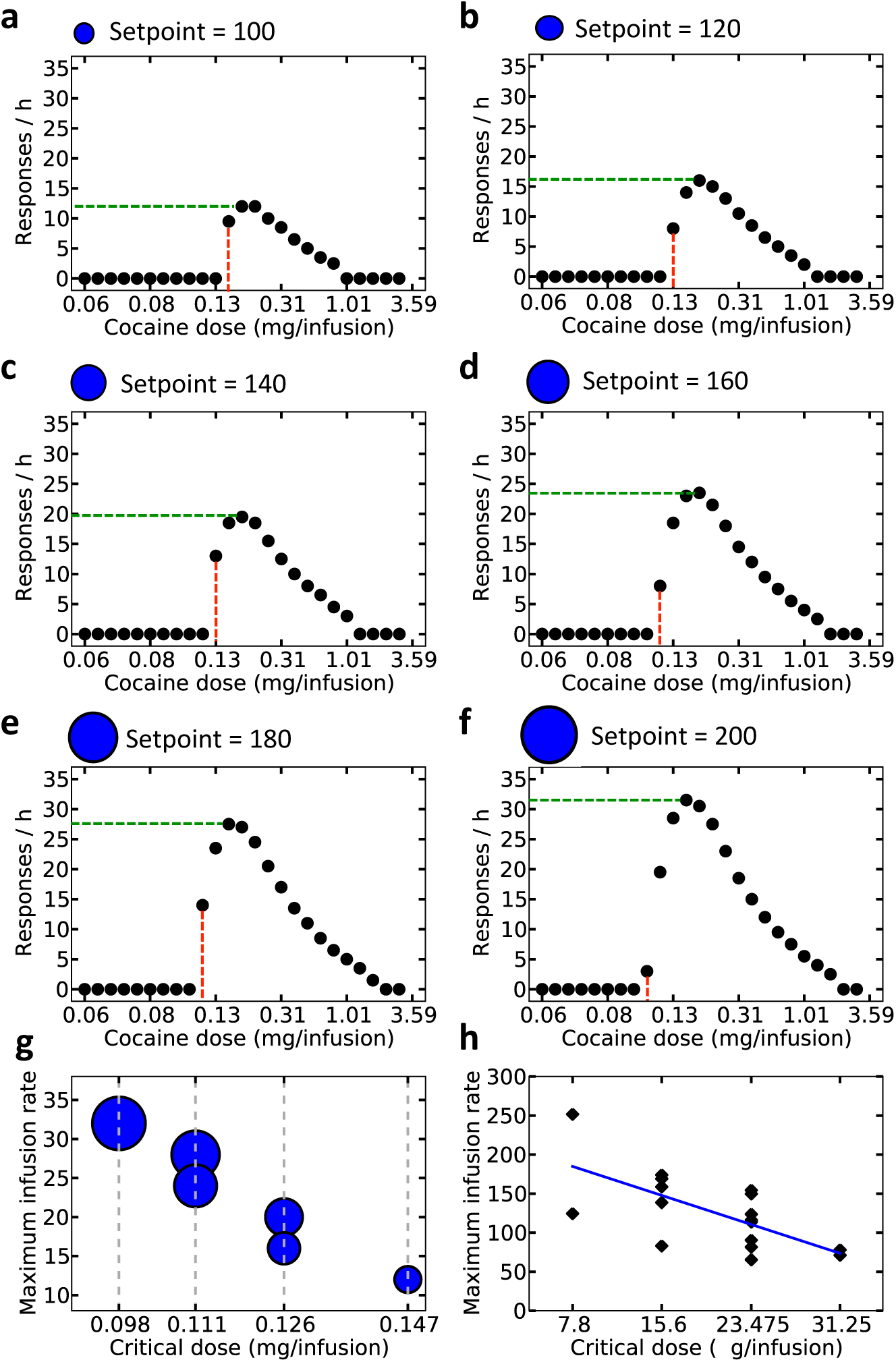
Simulation and experimental results on the interaction between critical unit dose and maximum infusion rate. Simulations show that as the setpoint level escalates (increasing order in panels A to F), the minimum unit dose (red line) at which the model shows motivation for seeking cocaine decreases, whereas the maximum infusion rate (green line) among all tested doses increases (simulation results summarized in panel G). (H) Experimental results of cocaine self-administration in rats (n=17) verified the negative correlation between these quantities (*p* < 0.01).

By further increasing the unit dose, the infusion rate steeply increases to a maximum level, due the increased drive-reduction reward of each unit dose and thus, increased motivation of drug seeking (). After reaching the maximum (dashed horizontal line in Fig. 8A-F), however, the infusion rate gradually decreases by further increasing the dose. This is because the combination of infusion rate and unit dose at this maximum point is just sufficient for reaching the setpoint and fluctuating around it. At higher doses, therefore, the agent should decrease the infusion rate in order to avoid overshooting (). Critically, our theory predicts that the higher the setpoint level is, the higher the maximum infusion rate among different unit doses will be (dashed horizontal line in Fig. 8A-F). This is again due to faster elimination of cocaine at higher rates, which requires higher infusion rate to compensate for it.

In sum, our theory proposes that the individual differences in dose-response curves (22) (see for experimental results) stem from differences in setpoint levels. Furthermore, it predicts that by increasing the setpoint level, the critical dose decreases, but the maximum infusion rate increases. This predicts a behaviorally-observable inverse correlation between the critical dose and the maximum infusion rate (Fig. 8G). To test this prediction, we analyzed previously published data from rats (n=17) self-administering different doses of cocaine (22), and revealed that this inverse correlation is in fact significant (*p* < 0.01) in experimental evidence (Fig. 8H).

## DISCUSSION

This study suggests that drug addiction is a pathological state of a goal-directed associative learning system that aims at defending the physiological stability of the organism. Critically, our theory is built upon a model-based RL system, which characterizes goal-directed planning (a.k.a, action-outcome associative structure) in animals (36). That is, the agent plans for seeking the outcome that fulfills its potentially escalating need for cocaine. This argues that many aspects of addiction, like compulsivity, that has been classically attributed to dominance of a habit system (1, 2) could be explained, along with many other evidence, by a goal-directed system.

Critically, habit-based theories of addiction fail at explaining several experimental results that are explained in this paper. The underlying assumption of those models (2, 4–6) is that drugs increase the phasic DA activity, which supposedly carries a reward prediction error signal (38). This drug-induced DA response would result in a maladaptive increase in the subjective value of drug-related choices. However, since higher doses of drug induce higher DA responses and thus higher reward prediction errors (20), those theories predict that increasing the unit-dose of drug should increase the animal’s motivation for drug self-administrations. Although this explains dose-dependent increase in the motivation to self-administer cocaine as measured in a progressive-ratio schedule, it is in contradiction with the well-known decreasing trend of the dose-response curve (13). Our theory explains both of these two seemingly contradictory behaviors. In our model, increasing the unit dose increases the drive-reduction rewarding effect of drug and thus, increases the breakpoint in a progressive ratio schedule. Increasing the dose, however, prolongs the post-injection satiation effect, resulting in longer inter-infusion intervals and thus, a decreasing trend in the dose-response curve (Fig. 3A).

Furthermore, habit-based theories leave the robust pattern of drug self-administration (13) (initial loading and pause phases, and the forthcoming regular responding) unexplained.

Despite the above argument, one could suggest a different way of implicating the habit system in addiction, that is, drugs hijack the homeostatic regulation system (the same way as proposed in this paper), but the generated rewards are learned by a habitual, rather than a goal-directed, system. Firstly, such an account is inconsistent with the fact that compulsive drug-seeking in human addicts is frequently goal-directed, as getting access to, procuring and taking drugs in the real world often require complex forward-looking behavioral strategies (39, 40). Devising such complex behavioral strategies is a clear indication of a goal-directed system being in control.

Secondly, such habit-based models fall short of explaining the regularities in the drug self-administration behavior (13) and the priming-induced relapse (19). The initial loading and pause phases followed by regular responding in the self-administration paradigm shows robust modulation of behavior by the internal state, which is inconsistent with the inflexible nature of habits. To explain those patterns, one could suggest adding an extra component to the habitual system for modulating the cached habit values in a generalized manner; that is, low/high levels of brain cocaine concentration energizes/ inhibits all responses, including the lever-press response. However, such an assumption is in contradiction with the pattern of priming-induced relapse (Fig. 6C, F). According to those data, higher levels of cocaine (i.e., bigger priming doses) increase press-lever, rather than decreasing it.

### Neural Substrates

Our model raises the important question of the neural parameters that are regulated during cocaine self-administration. At this stage, we can only speculate by integrating available, albeit incomplete, evidence. We postulate that the internal variable (*h_t_*) in our model is encoded by the relative excitability of direct- vs. indirect-pathway medium spiny neurons (MSNs) of the basal-ganglia (BG) to glutamatergic transmission from cortex; and that this relative excitability is modulated by striatal DA level through D1- and D2-like receptors in excitatory and inhibitory manners, respectively. Thus, as cocaine directly increases striatal dopamine (DA) level (26), it changes the internal state *h_t_*.

As evidence supporting this suggestion, it is demonstrated that during cocaine self-administration, rats first achieve and then maintain dopamine levels at an abnormally high level in the nucleus accumbens (41). This maintained level is significantly higher is LgA, as compared to ShA animals (41). This elevated cocaine-induced DA concentration in striatum, causing the rewarding effects of the drug (42), is transmitted to the downstream MSNs through two opposing channels: D1- and D2-like receptors. Whereas facilitation of D1R signaling strengthens the rewarding effects of cocaine (43), facilitation of D2R signaling attenuates it (44). Striatal DA level, in fact, enhances the excitability of D1R-expressing (D1R+), but decreases the excitability of D2R-expressing (D2R+) MSNs to cortical glutamatergic afferent. Thus, we hypothesize that cocaine reward is attained by increasing *h_t_*, defined by the relative excitability of D1R+ vs. D2R+ MSNs. Not only in drug context, but direct optogenetic activation of D1R+ and D2R+ MSNs is also shown to have rewarding and punishing effects, respectively (45), further supporting the hypothesis that increasing *h_t_* has reinforcing effects.

However, increasing *h_t_* (i.e., Δ*h_t_*) produces different rewarding effects at different initial levels of *h_t_*. Whereas D1R stimulation (equivalent to higher *h_t_*) attenuates motivation for cocaine seeking (i.e. motivation for increasing *h_t_*), D2R stimulation increases it (46). This indicates that the marginal reward induced by increasing *h_t_* gets smaller at higher level of *h_t_*. In other words, animals have less motivation for increasing *h_t_*, if *h_t_* is already high. Consistent with our model, this suggests existence of a setpoint, *h^*^*, against which *h_t_* is compared, and the animal’s motivation for cocaine depends on the extent to which its need (drive) is reduced by getting closer to *h^*^*.

Several lines of studies show long-lasting neuroplasticities in this circuit, due to chronic cocaine use. Chronic cocaine use in human addicts is associated with decreased D2R availability, increased density of DA transporters (30, 47), and decreased striatal DA release (32, 48). Similarly, long-term cocaine consumption induces decreased D2R availability in non-human primates (23, 31) and rats (49), and increased dendritic spine density in D1R+ MSNs in mice (50). The common consequence of all these adaptations is reduced effect of DA signaling on the downstream circuits, supposedly in order to compensate for their drug-induced over-activation. That is, chronic cocaine use lowers the ability of DA to increase the relative excitability of D1R+ compared to D2R+ MSNs. This is equivalent to our assumption that chronic cocaine elevates the setpoint level, *h^*^*.

According to our model, an escalated setpoint level results in higher homeostatic deviation under normal (i.e., non-drug) conditions, which in turn leads to a higher rewarding value (i.e., drive-reduction effect) of taking drugs (). This explains the inverse correlation between D2R availability and the reported pleasantness of taking psychostimulants, observed in human addicts (25)). Note that down-regulation of D2R is equivalent to elevation of the setpoint in our model.

D2R availability is also inversely correlated with motivation for drugs under drug conditions, as measured by steady-state rate of cocaine SA in monkeys (23) (). According to our model, lower D2R availability (i.e., escalated setpoint) motivates the animal to maintain striatal cocaine concentration at an elevated level (41). Due to faster elimination of cocaine at higher levels, however, defending homeostasis requires a higher infusion rate ().

### Testable Predictions

Our theory makes several predictions that are testable experimentally. As mentioned above, the model predicts that: (1) loading and pause phases of cocaine SA will be more pronounced by reducing the time-out period (); (2) there exist durations of access that are too short to induce escalation of cocaine intake but nevertheless sufficiently long to maintain it once developed (); (3) extinguishing, even fully, cocaine-conditioned interoceptive cues will not prevent relapse upon re-exposure to cocaine reinforcement (). (4) the response-triggering level of striatal cocaine (and DA) will be different during SA of different unit doses of cocaine (). (5) Last but not least, we expect the D2R level to be positively correlated with the critical unit dose at which animals start self-administering cocaine (G).

## Methods

### Simulated task

The model is simulated in an artificial environment where at each time point (every 4 seconds), the agent chooses between pressing an active lever, pressing an inactive lever, or doing nothing (representing grooming, sniffing, rearing, cocaine-induced stereotypy, etc.). Pressing the active lever results in an infusion of cocaine (fixed-ratio 1) delivered during 4sec, followed by a 20sec time-out during which pressing the lever has no consequence. Pressing the inactive lever is always without any consequence (see for the Markov Decision Process (MDP)). Pressing either of the two levers imposes a small cost to the agent, representing the energy spent for performing the response. As every action is assumed to take 4sec, simulating one day of an experiment is equivalent to 24*60*60/4 = 21600 trials (24 hours, every hour is 60 minutes, every minute is 60 seconds). During the rest of the day, both agents are in an environment where no action or state-transition is available. Therefore, the only variables of the model that might change during this period are the internal state (which returns back to zero at the beginning of this period, as the previously-administered cocaine degrades) and the homeostatic setpoint level (which gradually recovers back to the setpoint lower-bound).

During a pre-training period, the agent is simulated in the described MDP for 1hr per day, until the rate of cocaine infusion converges to a steady level. After this acquisition criterion was reached, LgA and ShA agents, as two instantiations of the pre-trained agent, have 6hr and 1hr daily access to the self-administration MDP, respectively. This MDP and the pre-training and training conditions explained above match the experimental tasks replicated in this paper, except when it is explicitly mentioned that a different MDP is used.

### State-space representation

The augmented state of the agent at each time is a mixture of its external (*s_t_*) and internal (*h_t_*) states; i.e. (*s_t_, h_t_*). The external state takes discrete values, whereas the internal state is a continuous variable. As the simplest way to tackle this inconsistency, we assume that the agent learns two separate representations of the MDP at the same time (): one for when the agent is totally under cocaine’s effect and the internal state is at the setpoint level (cocaine MDP), and another representation for when cocaine is absent and the internal state is at zero (no-cocaine MDP). The extent to which the agent exploits and updates each of these two representations depends on its internal state, *h_t_*. Variable *c_t_*, taking values between zero and one, defines the extent to which the agent is under the effect of cocaine:

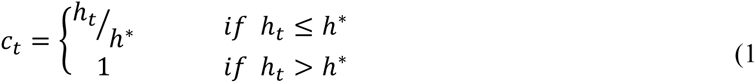

Accordingly, the augmented state-space of the agent can be defined by (*s_t_, c_t_*).

### Homeostatic mechanism

Upon each infusion of cocaine, the cocaine concentration rises rapidly and then eliminates during a longer time-course. The simulated pharmacokinetics of cocaine (and thus the internal state, Fig. 1B) is such that at each time unit, the agent absorbs 12% of the cocaine that is self-administered, but not absorbed yet. Also, irrespective of the choice made, 0.7% percent of the cocaine available in the brain degrades in every time unit. These two processes are equivalent to the following pharmacodynamics equation for the internal state:

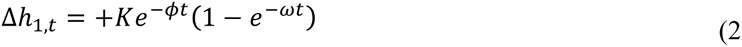

Where *ω* and *ϕ* are the absorption and elimination rates of cocaine, respectively. *K* is a constant that depends on the unit dose of cocaine. We set *K* = 50 for a single infusion of 0.250mg of cocaine. *K* changes proportionally for higher or lower unit doses. Clearly, repeated infusions results in the buildup of cocaine in the brain and thus, accumulation of drug influence on the internal state.

Note that an elevated homeostatic setpoint in LgA agents induces escalation because maintaining the internal state at a higher setpoint requires more infusions to compensate for the relatively faster elimination of cocaine when it has a high level in the brain. In fact, a constant elimination rate (0.7%) leads to higher total amount of elimination of cocaine at higher concentrations.

### Reward computation mechanism

As in the homeostatic RL theory (15), the drive level of the agent at each time, *t*, is computed as the distance (deviation) of the internal state, *H_t_*, from the homeostatic setpoint, *H^*^* (Fig. 1A):

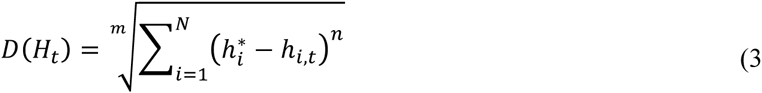

Where *N* is the number of regulated variables (dimensionality of the homeostatic space). For simplicity, we set *N* = 1 in the simulations, thus discount the interaction between the cocaine-related internal state and other regulated variables like glucose, temperature, etc.

Building upon the drive reduction theory of reward (15), we define the rewarding/punishing value of an infusion of cocaine as its ability to decrease/increase the drive level of the agent (15) (Fig. 1A):

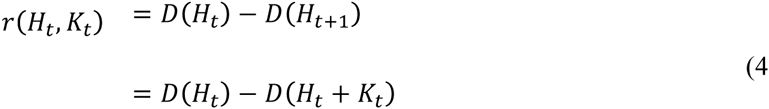

Where *K_t_* is the shift of the internal state upon cocaine infusion.

### Setpoint adaptation mechanism

We assume that the homeostatic setpoint undergoes slow adaptation is response to the strong effect of cocaine in the homeostatic regulation system. For simplicity, we assume a linear adaption, where the cocaine-related setpoint, *h\**, shifts forward in a magnitude proportional to the dose of cocaine, upon each infusion:

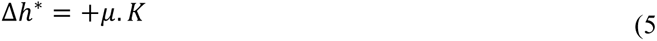
where *K* is the dose of cocaine, and *μ* is the adaptation rate. In every time point, the setpoint also undergoes a recovery to its normal level through an even slower adaptation process: 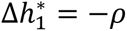.

Taking into account the neurobiological constraints on synaptic adaptation, we impose a lower bound and an upper bound on the setpoint, denoted by 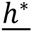 and 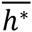 respectively (Fig. 1A).

### Learning mechanism

The model learns the environmental contingencies in the form of an outcome function, 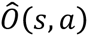, and a transition function, 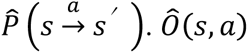 represents the expected outcome upon performing action *a* at state *s*. 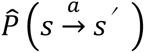 indicates the expected probability of arriving at state *s′*, upon performing action *a* at state *s*.

More precisely, as the internal state is also augmented into the state-space, the agent learns two separate outcome functions, 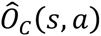 and 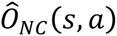, representing the expected outcome encoded in the cocaine and no-cocaine MDPs. Similarly, the agent learns two separate transition functions: 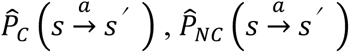. Supposing that the agent performs action *a* at state *s,* receives outcome *o* (dose of administered cocaine), and enters a new state *s′*, the outcome function will be updated as follows:

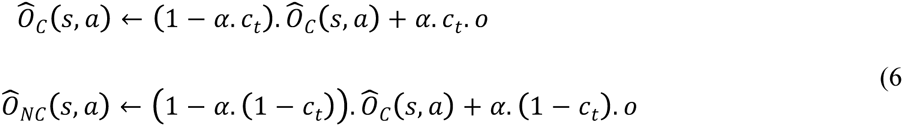
where 0 ≤ *α* ≤ 1 is the learning rate. In fact, the rate of update of each of the two outcome functions is proportional to the extent to which the agent is affected (*c_t_*) or not affected (1 − *c_t_*) by cocaine. Similarly, the transition functions will be updated as follow:

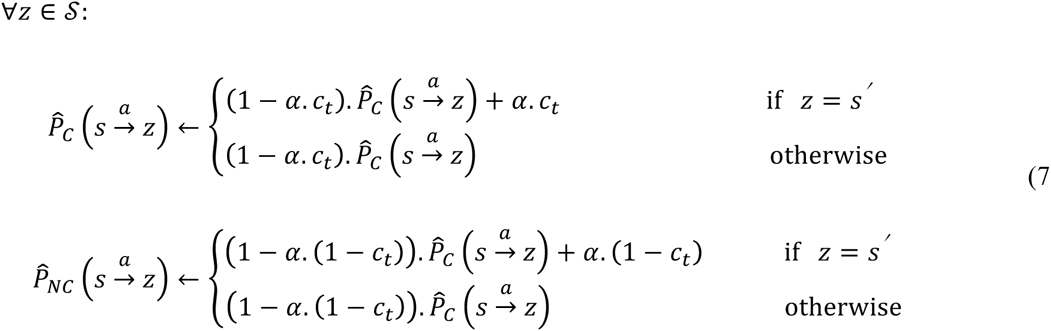

Where 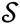 represent the set of all external states.

### Value estimation mechanism

The estimated value of each alternative is computed by doing a look-ahead goal-directed search based on the learned environmental contingencies (51):

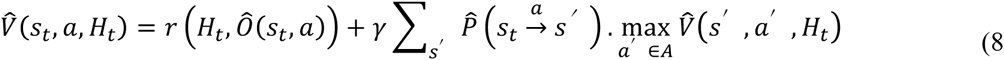
where 0 < *γ ≤* 1 is the discount factor. In fact, the rewarding value of a behavioral policy is equal to the sum of discounted rewards that the agent expects to receive by performing that strategy. The reward of each single action within this strategy is computed as the drive-reduction effect of the expected outcome of that action (equation 4).

More precisely, as the agent has learned two separate outcome and transition functions, using equation 8, the agent can compute two different value functions for the cocaine and non-cocaine 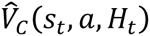 and 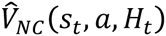. The weighted average of these two values is then used as the overall value assigned to a state-action pair:

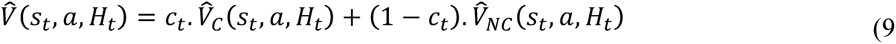

As before, 0 ≤ *c_t_ ≤* 1 represents the extent to which the agent is under the effect of cocaine.

Note that as the MDPs used in our simulation are cyclic, we limit the depth of tree search to three levels, in order to avoid infinite search time. Simulation results are not sensitive to the limit chosen for tree search.

### Action selection mechanism

Given its current state, *s_t_*, the agent chooses among possible options, with a probability proportional to their estimated values (*softmax* rule (51)):

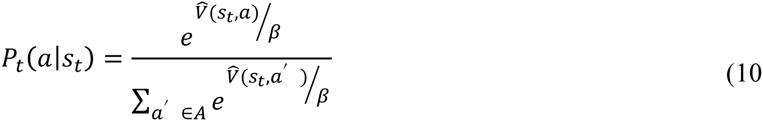
where 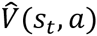 is the estimated value of taking action *a* from state *s_t_*, and *β* in the rate of exploration.

### Simulation details

The free parameters of the model and their values as used in the simulations are presented in. It is noteworthy that among all these parameters, the qualitative nature of simulation results is only sensitive to the values of *β* (rate of exploration) and electric-shock punishment (only present in Fig. 5). Other free parameters can take0020a wide range of values, without affecting the essential behavior of the model.

All the results (except for results in Fig. 5) are derived from simulating each agent only once (rather than averaging over several simulations). However, the results are robust, since the only source of stochasticity in the model is exploration (due to using the softmax action-selection rule). However, in simulation results, this stochasticity is averaged out over the several hours of simulated cocaine self-administration by the same agent.

### Experimental methods

#### Subjects

A total of 19 5-month old, male Wistar rats (Charles River, L’Arbresle, France) were used in this experiment. They were housed in groups of 2 and were maintained in a light- (reverse light-dark cycle) and temperature-controlled vivarium (21±2°C). All behavioral testing occurred during the dark phase of the light-dark cycle. Food and water were freely available in the home cages throughout the duration of the experiment. Home cages were enriched with a nylon gnawing bone and a cardboard tunnel (Plexx BV, The Netherlands).

#### Ethical statement

The experiment was carried out in accordance with institutional and international standards of care and use of laboratory animals [UK Animals (Scientific Procedures) Act, 1986; and associated guidelines; the European Communities Council Directive (2010/63/UE, 22 September 2010) and the French Directives concerning the use of laboratory animals (décret 2013-118, 1 February 2013)]. The animal facility has been approved by the Committee of the Veterinary Services Gironde, agreement number A33-063-922.

#### Training history

All rats were selected from a previous experiment where they were extensively trained under a fixed-ratio (FR) schedule of reinforcement to press a lever to self-administer cocaine (0.25 mg per injection) through an indwelling intravenous catheter. Other general procedural information concerning intravenous cocaine self-administration (e.g., intravenous surgery; operant chambers) can be found elsewhere(19). Rats were trained during 38 daily sessions of 3 h and, as a result, self-administered a total amount of 209.5 ± 26.1 mg of cocaine per rat before being tested under the within-session dose shift procedure.

#### Within-session dose shift procedure

This procedure was designed to measure how rats adjust their rate of cocaine self-administration to a large, non-signaled change in unit dose within a session. Rats had first access to a high dose of cocaine (1 mg per injection) during the first 2 h of the session and then to a much lower dose of cocaine (0.0625 mg per injection) during the rest of the session. No signal announced this large within-session decrease (i.e., 16 fold) in unit dose. In addition, to avoid any limitation on the rate of cocaine self-administration, particularly at the low dose, no programmed time-out period followed the injections. Finally, to further incite rats to pay attention to the interoceptive effects of cocaine, no programmed response-contingent cue signaled drug reinforcement. Rats were tested under this procedure for a total of 8 daily sessions of 3–4h until stabilization of behavior (i.e., no ascending or descending trend in performance over 3 consecutive sessions). Only data obtained during the last 3 stable sessions are presented and analyzed here.

### Data analysis

The inter-infusion intervals (III) for the 19 rats, during the last three sessions of the experiment were analyzed. Every point on Fig. 7i is the average III over 19 rats and over the last three trials (averaged over 3*19=57 IIIs), for the *n*-th III after reducing the unit-dose of cocaine. To test the statistical significance of the IIIs being lower in the initial post dose-reduction self-administrations, we assume that III becomes stable after 15 post-reduction responses. Thus, for each animal and for each of the three sessions, we can use the IIIs between the 15^th^ and the 25^th^ post-reduction responses as the baseline III for that animal and that session (25 is the minimum number of post-reduction responses among all animals; i.e., the number of responses that all animals achieved). We then compare each individual IIIs with this baseline III. That is, we compute the difference between the first post-reduction III for each rat in each session, with every IIIs that is between the 15^th^ and 25^th^ post-reduction response, for that rat in that session. This gives 10 data points for each rat in each session. We pool together all these 10-data-points sets from all animals, from all three sessions, providing us with the total number of 10*19*3=570 data points. If these points are statistically greater than zero, it means that the first post-reduction III is smaller than the baseline III. The first point in Fig. 7j represents the *p-value* of such test (one-sided t-test). The next point shows the *p-value* of the same analysis, but the baseline III is compared to the “second” post-reduction III; and so on.

## AUTHOR CONTRIBUTIONS

M.K., B.G. and S.H.A. defined the project. M.K. designed the model, performed the simulations and analyzed experimental data. P.G. and A.D. performed the experiment. S.H.A. provided the raw data from previous experiments. M.K., S.H.A. and B.G. discussed the results. M.K. wrote the paper with the help of S.H.A. and B.G. All authors critically reviewed content and approved final version for publication.

## ACKNOWLEDGEMENTS

We thank Peter Dayan for critical discussions. M.K. and B.G. are supported by grants from Frontiers du Vivant, the French MESR, CNRS, INSERM, ANR, ENP and NERF. S.H.A. is supported by the French Research Council (CNRS), the Université de Bordeaux, the French National Agency (ANR, ANR2010-BLAN-1404–01), the Conseil Régional d’Aquitaine (CRA20101301022; CRA11004375/11004699) and the LabEX BRAIN (ANR-10-LABX-43).

## Supplemental Information

### Supplementary Figures

**Fig. S1.**
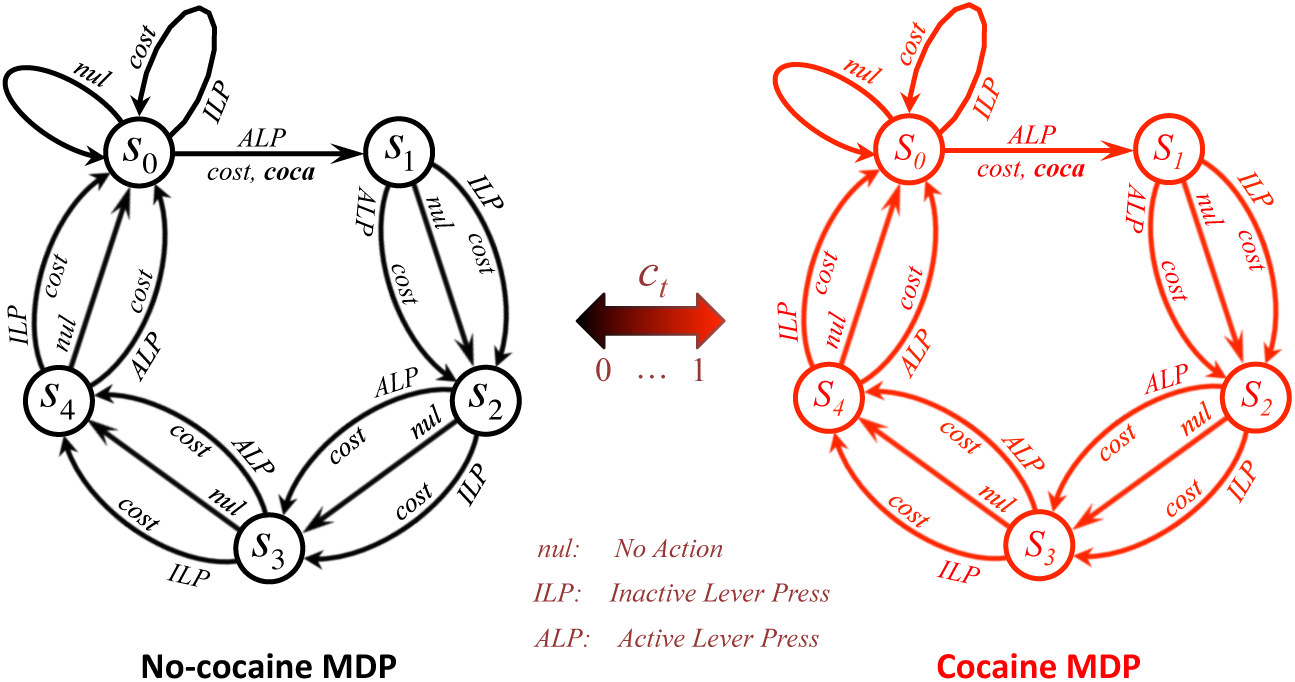
The Markov Decision Process (MDP) of the simulated cocaine-seeking task, in a fixed-ratio one (FR1) schedule, with 20 seconds time-out. Starting at state *s*_0_, the agent can press the active lever (*ALP*) to receive a certain dose of cocaine. In other states, *ALP* does not result in cocaine. Every action (arrow) is supposed to take 4 seconds to be performed. Therefore, starting from state *s*_0_ and pressing the active-lever, it will take 5*4 = 20 seconds to return to the initial state (*s*_0_) where cocaine can be self-administered again. Pressing the inactive lever (*ILP*) or doing nothing (*null*) has no consequence. Pressing either the active or the inactive lever has a fixed cost, representing the energy spent for performing such actions. The subjective representation of the MDP consists of two parallel MDPs, one for representing the world when the agent is under cocaine (cocaine MDP), and another one for representing the world when the internal level is at zero (no-cocaine MDP). The extent to which these two MDPs are exploited or updated depend on the extent to which the agent is under cocaine.

**Fig. S2.**
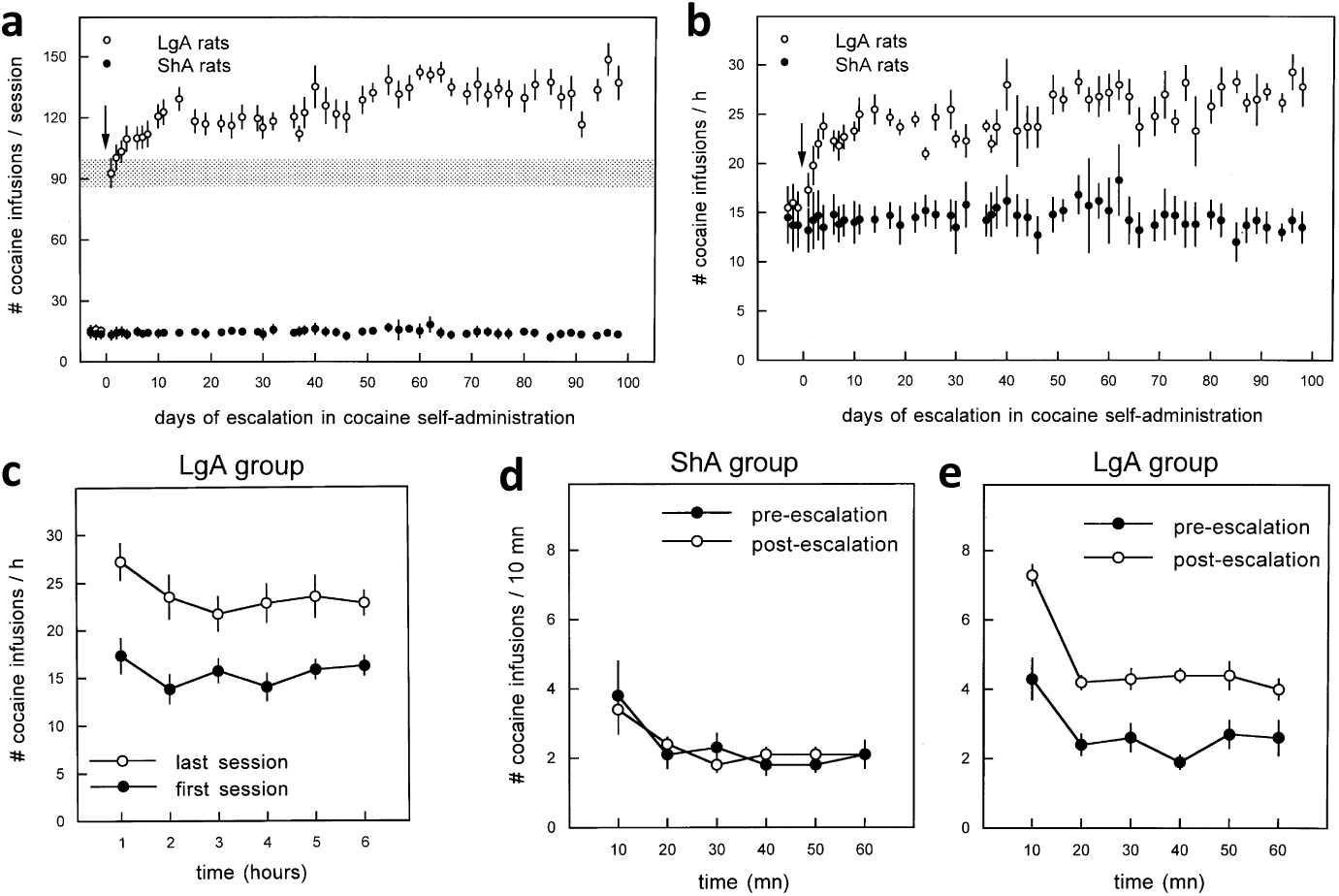
Experimental results (Ahmed and Koob, 1999) on the effect of the duration of drug-choice availability on the rate of consumption. Whereas the number of infusions per session remains stable in the ShA rats, it escalates in LgA animals (a). The same is true about the rate of infusion during the first hour of sessions (number of infusions/hr) (b). Similarly, the rate of infusion in intervals of 10min does not change from the first to the last session, in the ShA group (d), whereas a significant escalation is observed in the LgA group (c and e). Also, in both LgA and ShA rats, and during both pre- and post-escalation sessions, the rate of infusion is higher during the first 1 Omin interval, compared to the five next intervals (d and e). This phenomenon is known as “loading effect”.

**Fig. S3.**
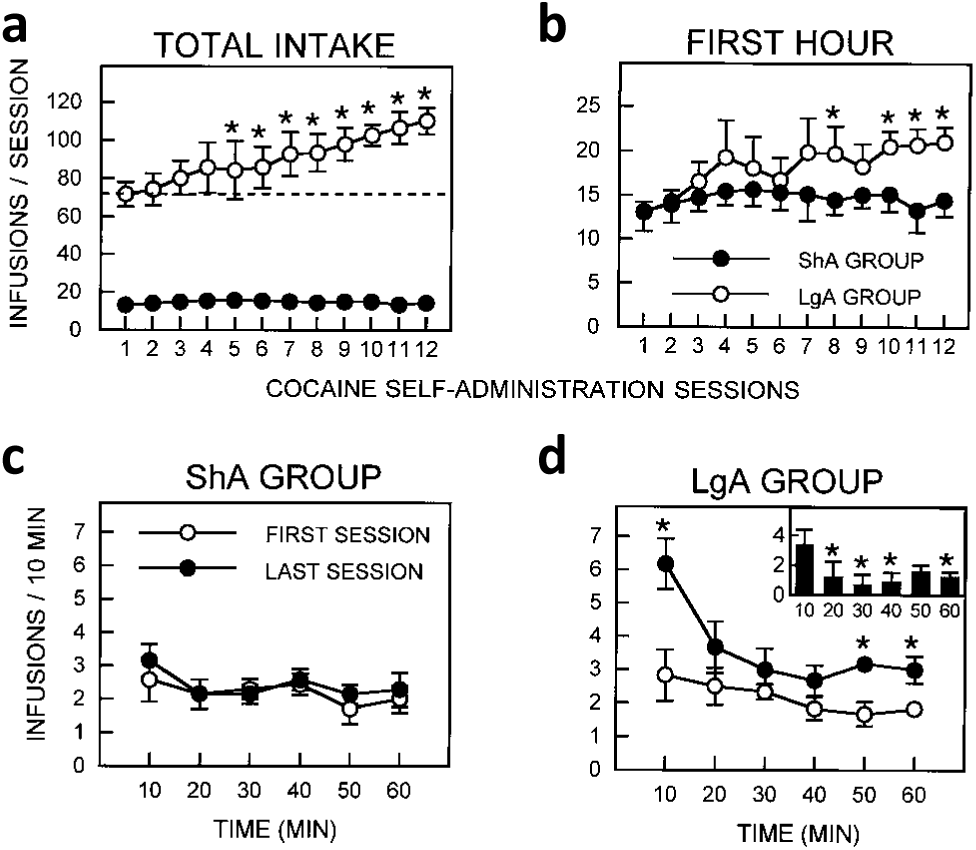
Experimental results (Ahmed and Koob, 1998) on the effect of the duration of drug-choice availability on the rate of consumption. Whereas the number of infusions per session remains stable in the ShA rats, it escalates in LgA animals (a). The same is true about the rate of infusion during the first hour of sessions (number of infusions/hr) (b). Similarly, the rate of infusion in intervals of 10min does not change from the first to the last session, in the ShA group (c), whereas a significant escalation is observed in the LgA group (d). This escalation, as shown in the inset in panel d (last minus first session), is stronger during the first 10min interval. Also, the rate of infusion in the last session of the LgA animals is higher during the first 10min interval, compared to the five next intervals. This phenomenon is known as “loading effect”. Whereas in this study (Ahmed and Koob, 1998) the loading effect is significant only in the last session of the LgA group, later studies show significant loading also in ShA animals, as well as in the first session (Ahmed and Koob, 1999).

**Fig. S4.**
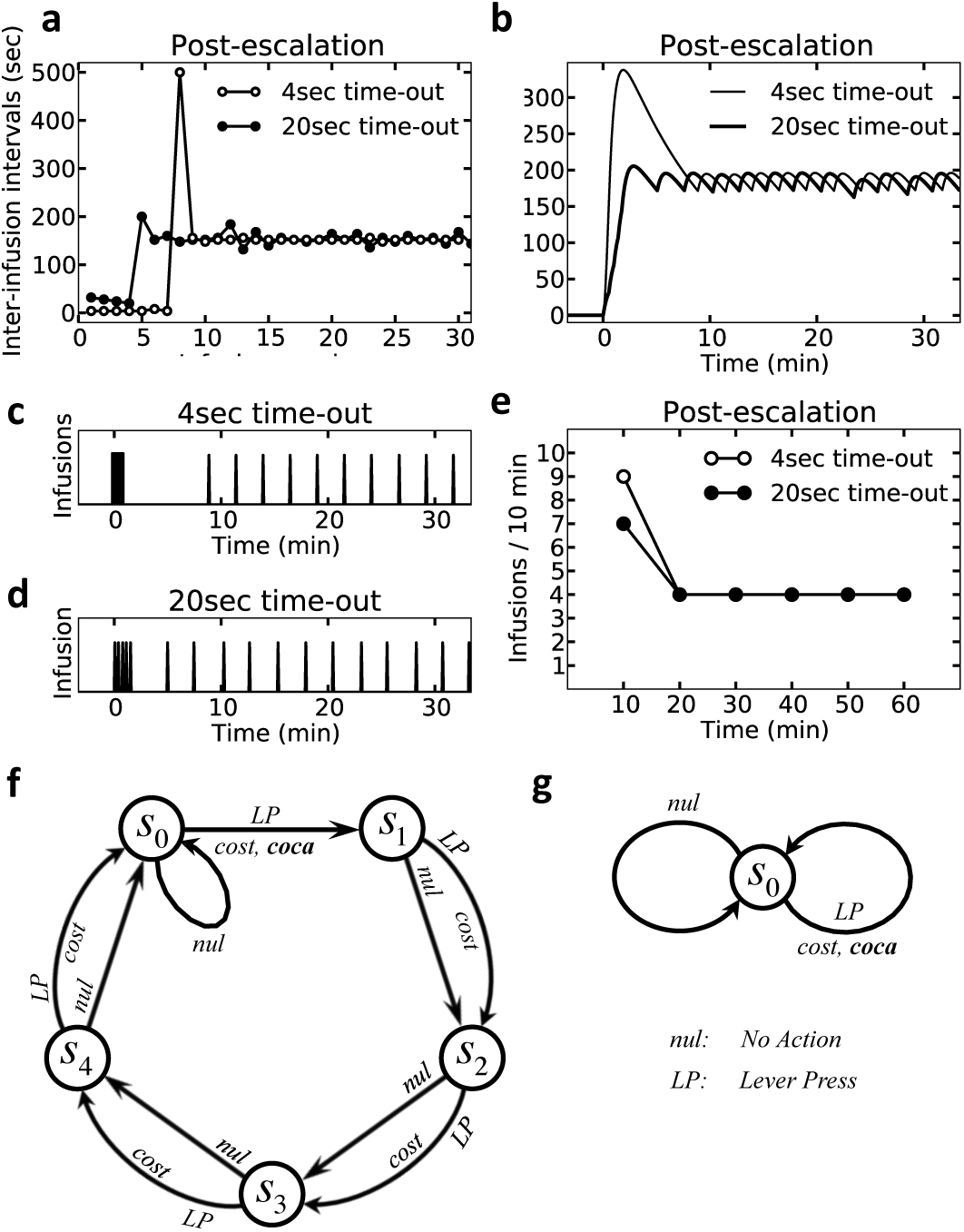
Simulation results predicting more pronounced load and pause effects in the 4sec, compared to the 20sec time-out task. In both 4sec and 20sec cases, the agents start the session in a cocaine-deprived state. Thus, they self-administer cocaine several times, with the lowest possible inter-infusion intervals (a). This period is known as loading phase. However, due to the pharmacodynamics of cocaine, after each infusion, it takes several seconds before cocaine reaches its maximal effect on the internal state (Fig. 1b). In the 20sec case, as the time-out is relatively long, the effect of every cocaine infusion on the internal state is almost completely applied before the next self-administration becomes available. In this condition, the agent’s internal state reaches the setpoint after five infusions (c). After this loading phase, the agent self-administers steadily. In the 4sec case, however, even though the first few infusions are sufficient for reaching the setpoint, their effect arrives much later than when self-administration is available again. Thus, the agent continues taking cocaine (a, b) for several extra times. These extra infusions result in overshooting the setpoint (d) after their effect on the internal state arrives. In order to return to the setpoint, the agent pauses taking cocaine for several minutes (b), resulting in one significantly large inter-infusion interval (b), known as the pause effect. After that, the agent self-administers steadily. Therefore, the model predicts that both loading and pause phenomena will more pronounced by decreasing the time-out duration. As in our model the circulating cocaine degrades faster when it is at higher levels, the overshoot of cocaine level in the 4sec case results in more cocaine elimination. In order to compensate for that, the agent takes more infusions of cocaine. As a result, the rate of infusion in the first ten minutes is higher for the 4sec case, than for the 20sec case (e). Plots f and g show the Markov Decision Process used for simulating the 20sec and 4sec cases, respectively.

**Fig. S5.**
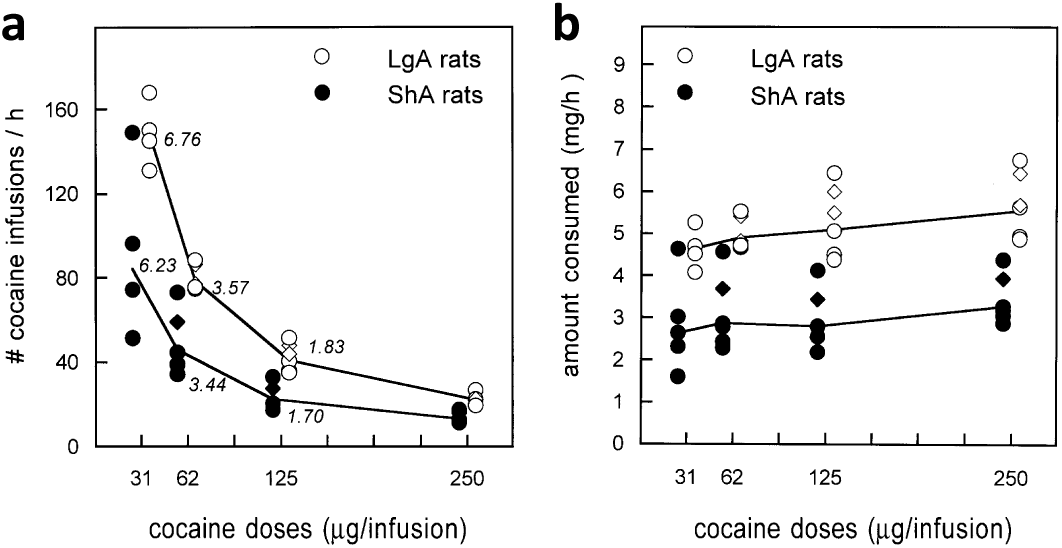
Experimental result (Ahmed and Koob, 1999) on post-escalation dose effect on infusion rate and total intake of cocaine. After 22 sessions of cacaine self-administration(250 (μg/injection) under LgA and ShA conditions, the rate of infusion was measured during four different sessions, for four different unit doses of cocaine: 31.25, 62.5, 125 and 250 μg. These sessions were dispersed between sessions 22 and 44, in a random order for each animal. As the dose increases, the rate of infusion decreases (a). For all doses, the rate of infusion is higher for LgA, compared to ShA rats. The total amount of consumption over one hour shows no significant change at different doses (b).

**Fig. S6.**
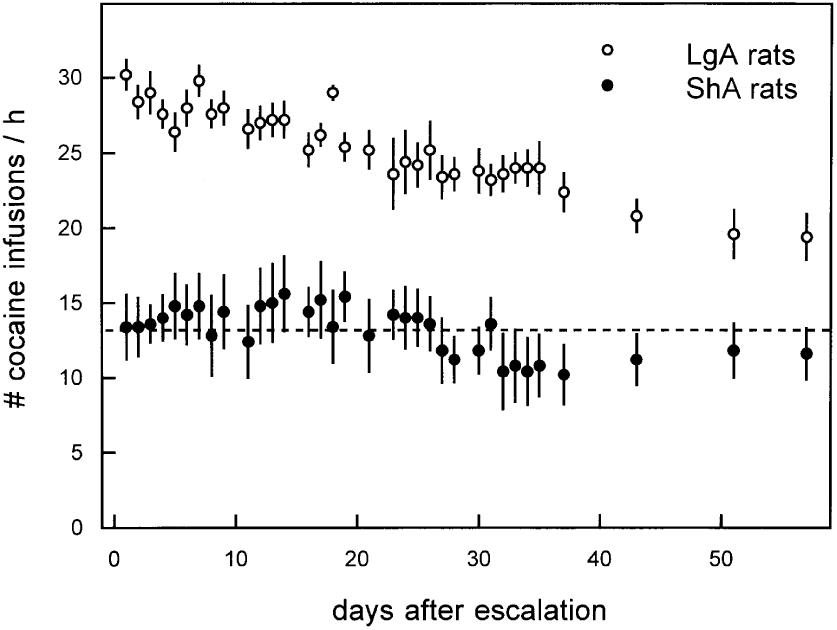
Experimental results (Ahmed and Koob, 1999) on post-escalation reduced availability of cocaine. After escalation (as in (Fig. S2), both LgA and ShA animals were given 1hr access to cocaine self-administration. Of a total number of 34 post-escalation sessions, the first 31 sessions were performed 5–6 days pre week, and sessions 32, 33, and 34 were performed once per week. Whereas the rate of infusion remained stable for the ShA rats, it decreased gradually in the LgA group.

**Fig. S7.**
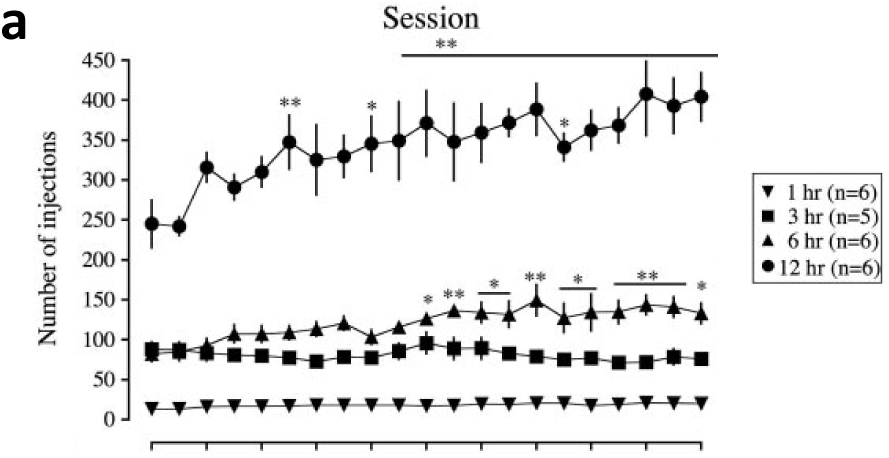
Experimental results (Wee et al., 2007) on the effect of session-duration on escalation. Results show that cocaine self-administration increased under 6hr and 12hr access conditions, but not under 1hr and 3hr conditions. The escalation of infusion rate was faster under 12hr, compared to the 6hr condition.

**Fig. S8.**
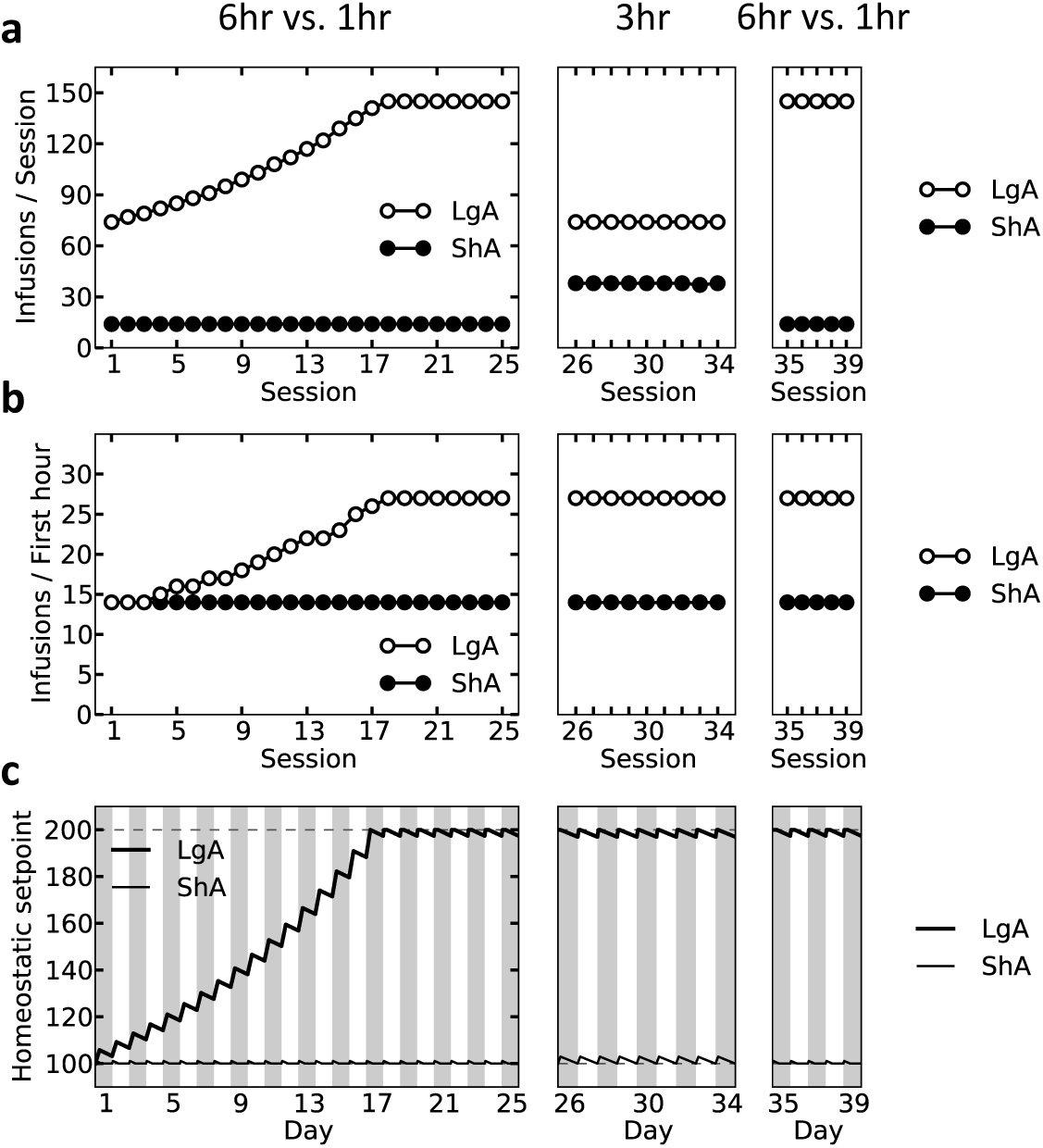
Simulation results predicting that 3hr access does not induce escalation, but keeps the escalated animals at an escalated level. After 25 sessions of 6hr vs. 1hr access (left panels), both LgA and ShA agents were given 3hr access per day, for 9 consecutive days. The elevation of the setpoint during 3hr is virtually equal to its recovery during the rest of the day (21hr). As these two processes cancel out each other, the setpoint level remains steady under 3hr access condition (plot c, middle). As a result, the rate of infusion/hr remains constant for both ShA and LgA agents (plots a and b, middle). Thus, if after the 3hr access phase, the agents return beck to the 6hr vs. 1hr access conditions, their infusion rate will be equal to the initial steady-state level (panels a and b, right).

**Fig. S9.**
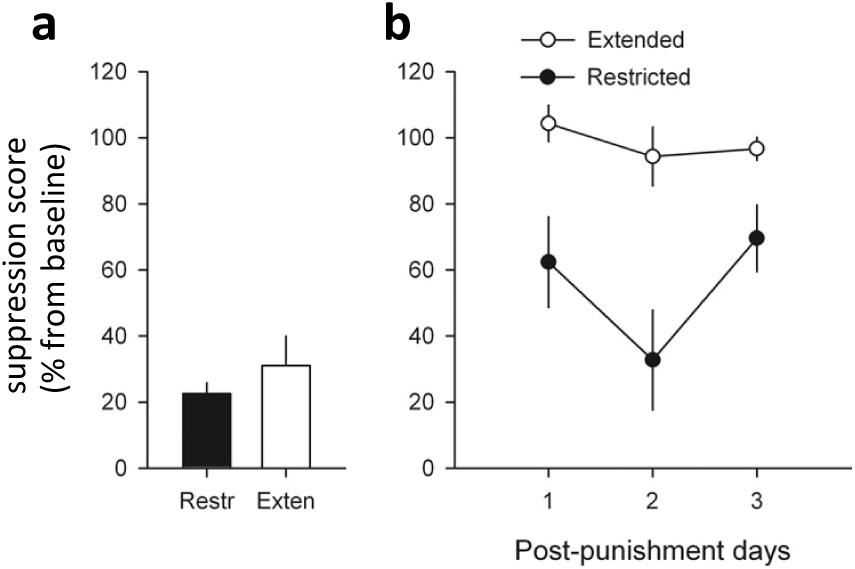
Experimental results (Ahmed, 2012) on the effect of extended drug-access on punishment-induced suppression of cocaine SA. Following 78 sessions of differential access to cocaine, both ShA and LgA had access to cocaine for 45 minutes during which, cocaine infusion was paired with an electric shock. Both groups decreased the rate of self-administration when cocaine infusion was paired with punishment (a). During the post-punishment days (three 45-minute sessions), LgA rats resumed self-administration more rapidly than ShA animal, which refrained from self-administering cocaine during at least three consecutive days (b). Exten: LgA condition; Restr: ShA condition.

**Fig. S10.**
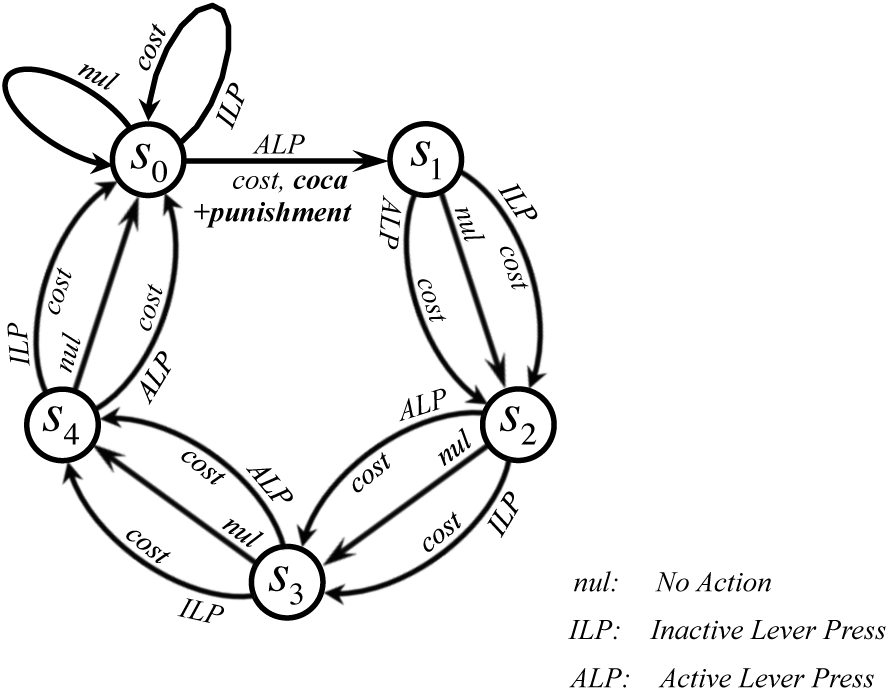
The Markov Decision Process (MDP) used for simulating the effect of extended drug-access on punishment-induced suppression of cocaine self-administration (Ahmed, 2012). A strong punishment (negative value, see Table. S1) was assigned to the same action that results in cocaine infusion. We simulate 12 instantiations of the model, 6 under LgA and 6 under ShA conditions. This is equal to the number of rats used in the corresponding experimental study (Ahmed, 2012). In both ShA and LgA groups, all agents equally decreased the level of self-administration, when cocaine infusion was paired with a strong punishment during a 45-minute session (Fig. 5a). During the post-punishment period, all the six LgA agents rapidly resumed the rate of infusion at the baseline level (Fig. 5b). This is because in spite of the expected punishment, the estimated value of lever-press was still high enough (due to the high value of cocaine) to motivate exploration of the lever-press action. Sufficient explorations of this option in the new punishment-free condition results in resumption of cocaine self-administration. In ShA agents, however, the rewarding value of cocaine is relatively lower (due to non-escalated setpoint). Thus, being paired with a strong punishment reduces the value of the lever-press action to such a low level that even when the punishment is removed, the agents explore this action extremely rarely. Therefore, the chance of updating the subjective representation of punishment probability is relatively lower in ShA agents, which results in delayed resumption of cocaine self-administration (Fig. 5b).

**Fig. S11.**
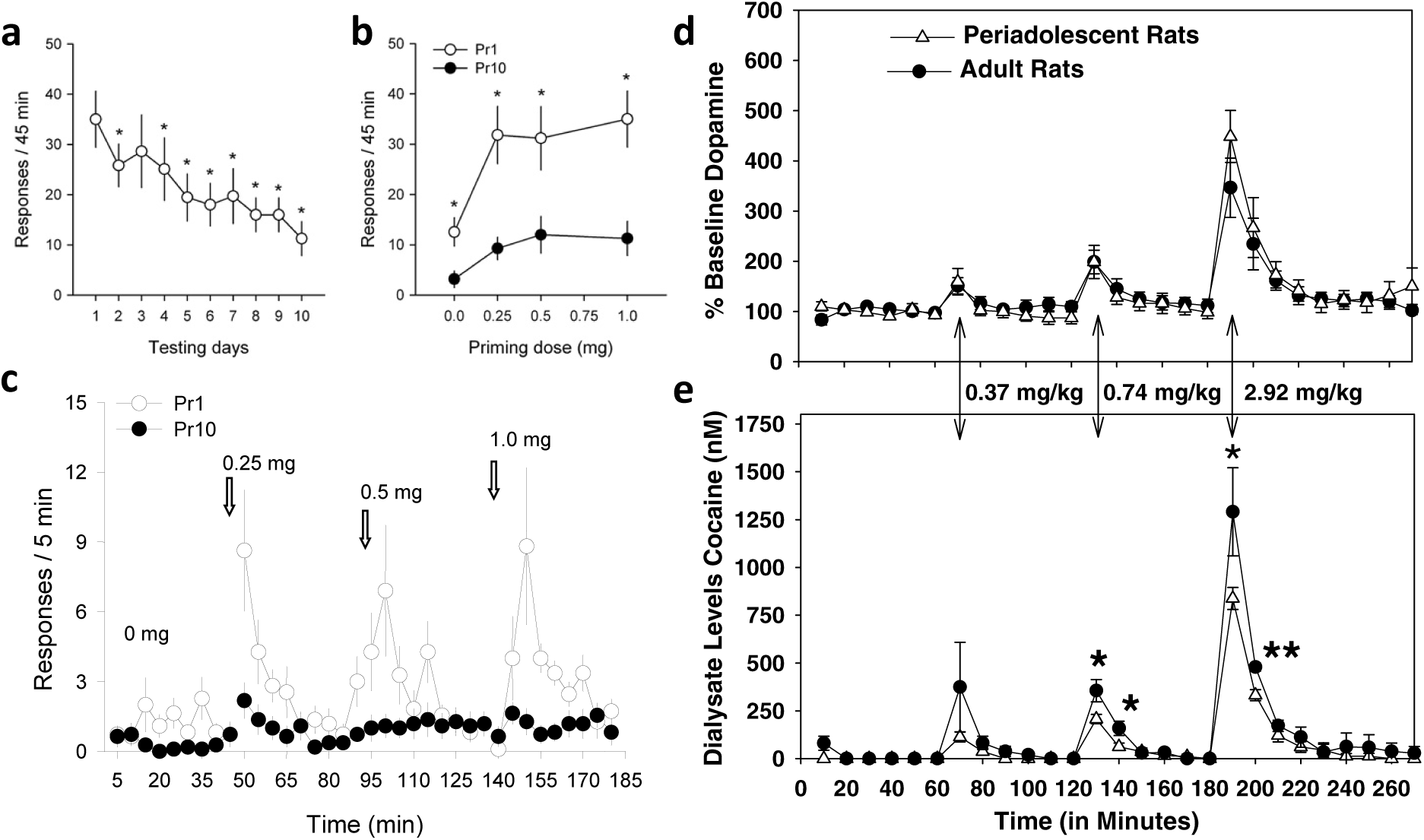
Experimental results (Frantz et al., 2007; Mihindou et al., 2011) on the extinction of priming-induced reinstatement. Following 32 sessions of 6hr access to cocaine, animals underwent a priming-induced reinstatement procedure during 10 consecutive days. Each day consisted of 5 consecutive sessions of 45min during which pressing the lever had no consequence (extinction). At the beginning of each session, the agent received a single priming injection of cocaine with the following doses: 0, 0, 0.25, 0.5, and 1mg. The rate of lever-press in the 45min upon infusion of the highest dose (1mg) decreased progressively (a) representing a gradual extinction of relapse. Such extinction also happened for other dose of cocaine, by comparing the first (Pr1) and the last (Pr10) extinction sessions (b). Zooming on response rates at 5min intervals showed a more precise pattern of priming induced reinstatement (c). On the first day of extinction, responding increased instantaneously upon injection of the lowest dose (0.25mg). For higher doses, however, the peak of response rate was achieved in the second (for 0.5mg) and third (for 1mg) 5min intervals. Plots d and e show the measured levels of dopamine and cocaine in the brain, respectively. Panels a, b, and c are reprinted from (Mihindou et al., 2011). Panels d and e are reprinted from (Frantz et al., 2007).

**Fig. S12.**
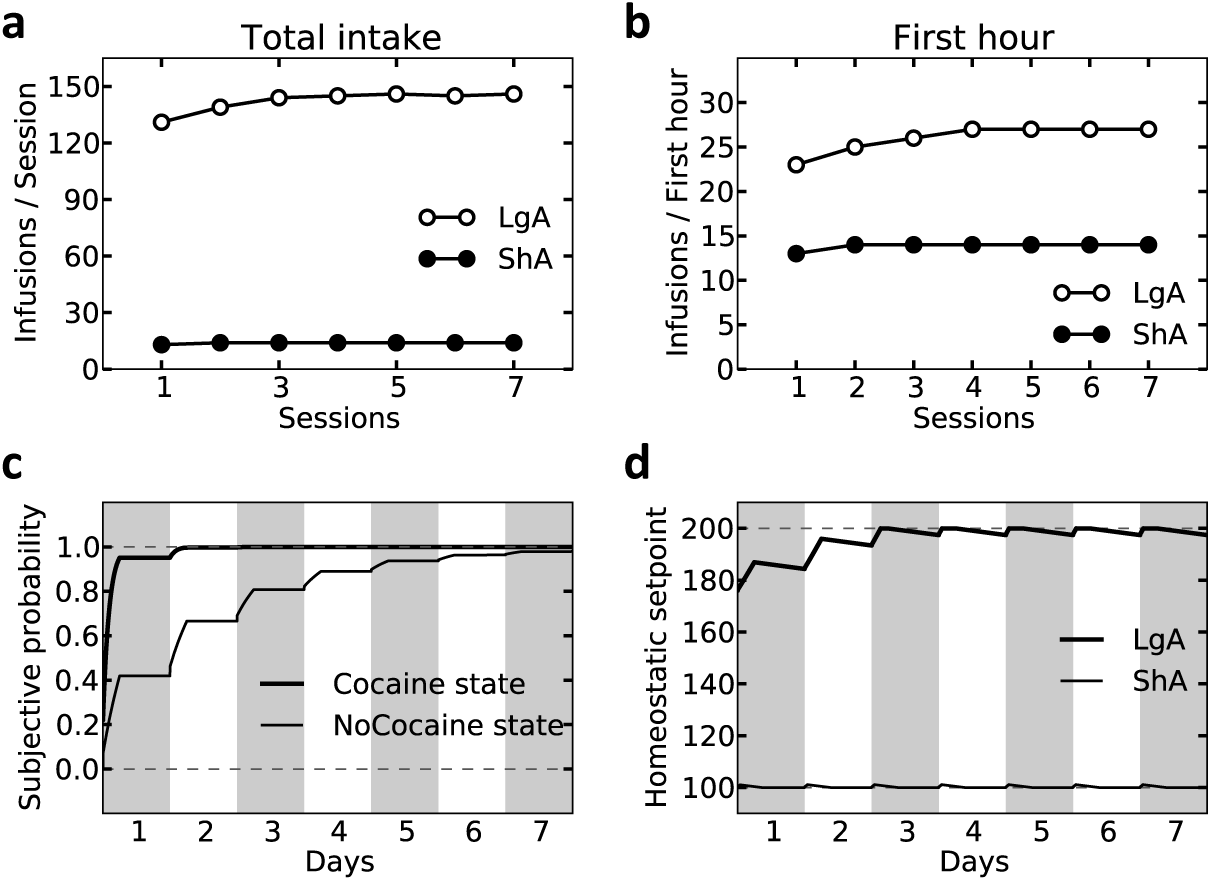
Simulation results predicting rapid re-escalation of cocaine SA, after extinction of priming-induced relapse. LgA and ShA agent underwent 25 sessions of 6hr vs. 1hr of cocaine self-administration, respectively. They then experienced 10 days in the relapse-extinction schedule as in Fig. 6. After this phase, the agents were again given 7 days of 6hr vs. 1hr access to cocaine self-administration. The rate of infusion by the LgA agent was at an escalated level, even in the first session of re-escalation (a, b). This is because the setpoint level was still at an elevated level (d). In fact, the 10-day extinction phase did not lead to recovery of the setpoint (Fig. 6d), and the extinction of relapse was only due to decreased subjective probability of receiving cocaine (Fig. 6e). As the subjective probability can be re-learned rapidly within the first session of re-escalation (c), and as the setpoint is still at a high level (d), cocaine infusion rate re-escalates rapidly after extinction of drug-induced reinstatement.

**Fig. S13.**
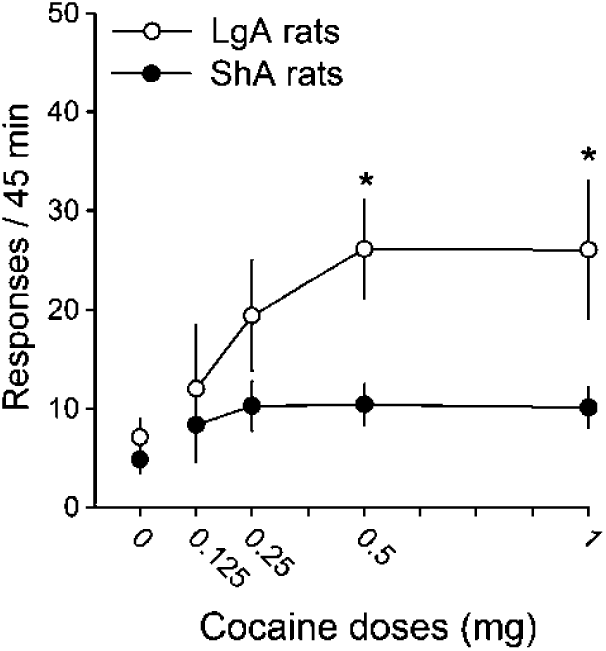
Experimental results (Ahmed and Cador, 2006) on the effect of extended drug-access on dose-dependent priming-induced reinstatement. Following 32 sessions of differential access to cocaine, both ShA and LgA agents passively received increasing intravenous doses of cocaine, one dose every 45 min with the first 45-min interval corresponding to behavioral extinction. During reinstatement testing, pressing tine lever had no consequence. Priming-induced reinstatement was pronounced more significantly in LgA, than in ShA rats.

**Fig. S14.**
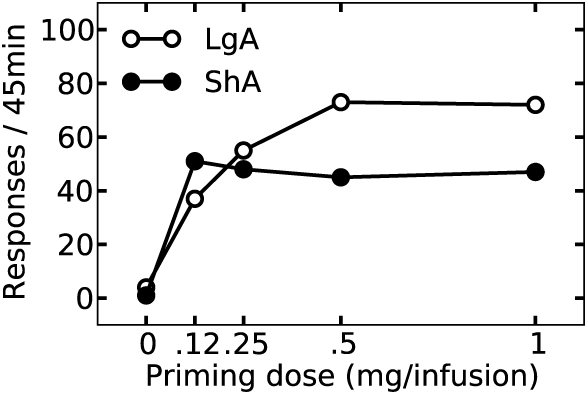
Simulation results replicating experimental data (Ahmed and Cador, 2006) (Fig. S13) on the effect of extended drug-access on dose-dependent priming-induced reinstatement. After 25 days of 6hr vs. 1hr access to cocaine, both LgA and ShA agents were provided with one 45-minute session in which, pressing the lever had no consequence. This extinction session was followed by five additional 45-minute extinction sessions, at the beginning of each of which, the agents received different priming doses of cocaine (0, 0.125, 0.25, 0.5, and 1mg). Priming-induced reinstatement was more pronounced in the LgA agent.

**Fig. S15.**
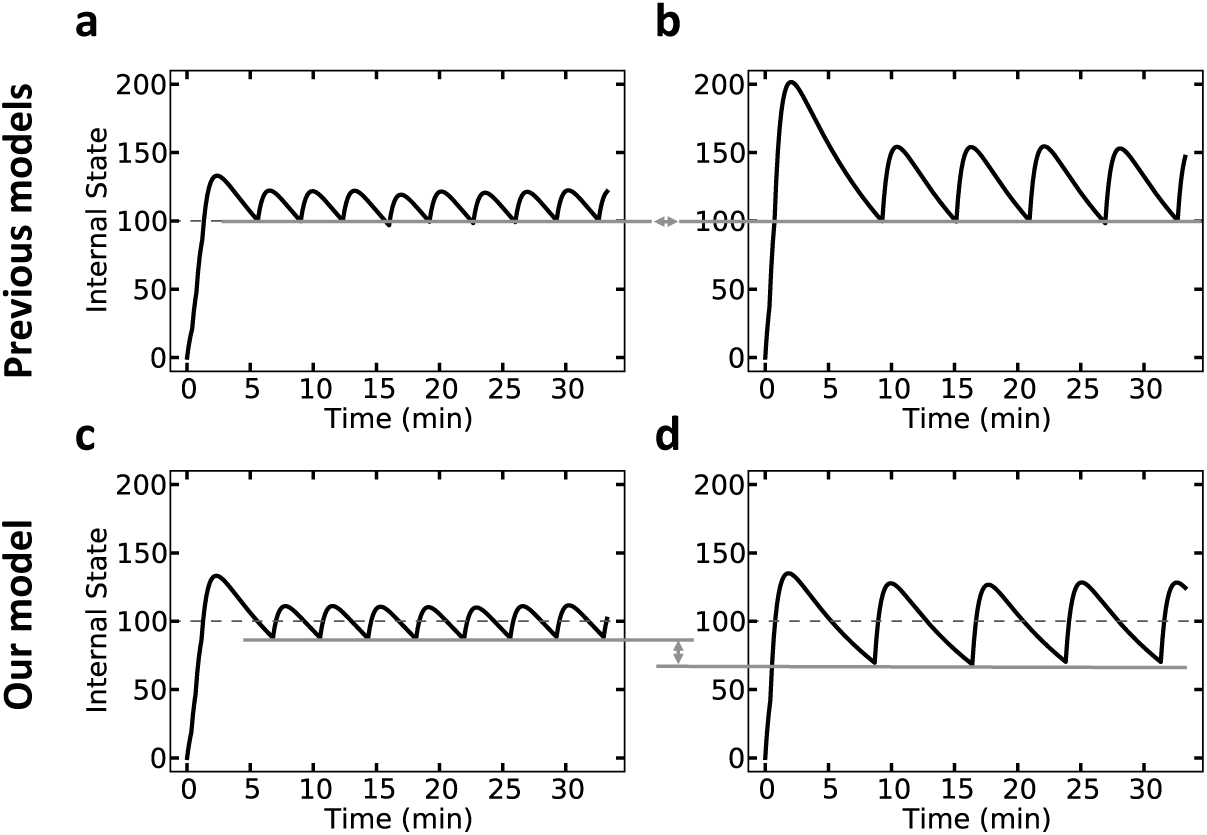
Simulation results predicting different satiety thresholds for different doses of cocaine. According to the previous models of cocaine self-administration, cocaine-taking response is generated by the animal as the internal state drops below a certain threshold (Ahmed and Koob, 2005; Tsibulsky and Norman, 1999) (a, b). Accordingly, these models predict that the lower bound of the cocaine level in the brain is equal, for all doses of self-administered cocaine (compare plots a and b). In our model, however, the agent’s objective is to keep its internal state as close as possible to the homeostatic setpoint (the setpoint is shown by a dashed line in plots c and d). Thus, our model predicts that the lower bound of cocaine level for a low dose of cocaine (c) will be higher than that of a high dose of self-administered cocaine (d).

**Fig. S16.**
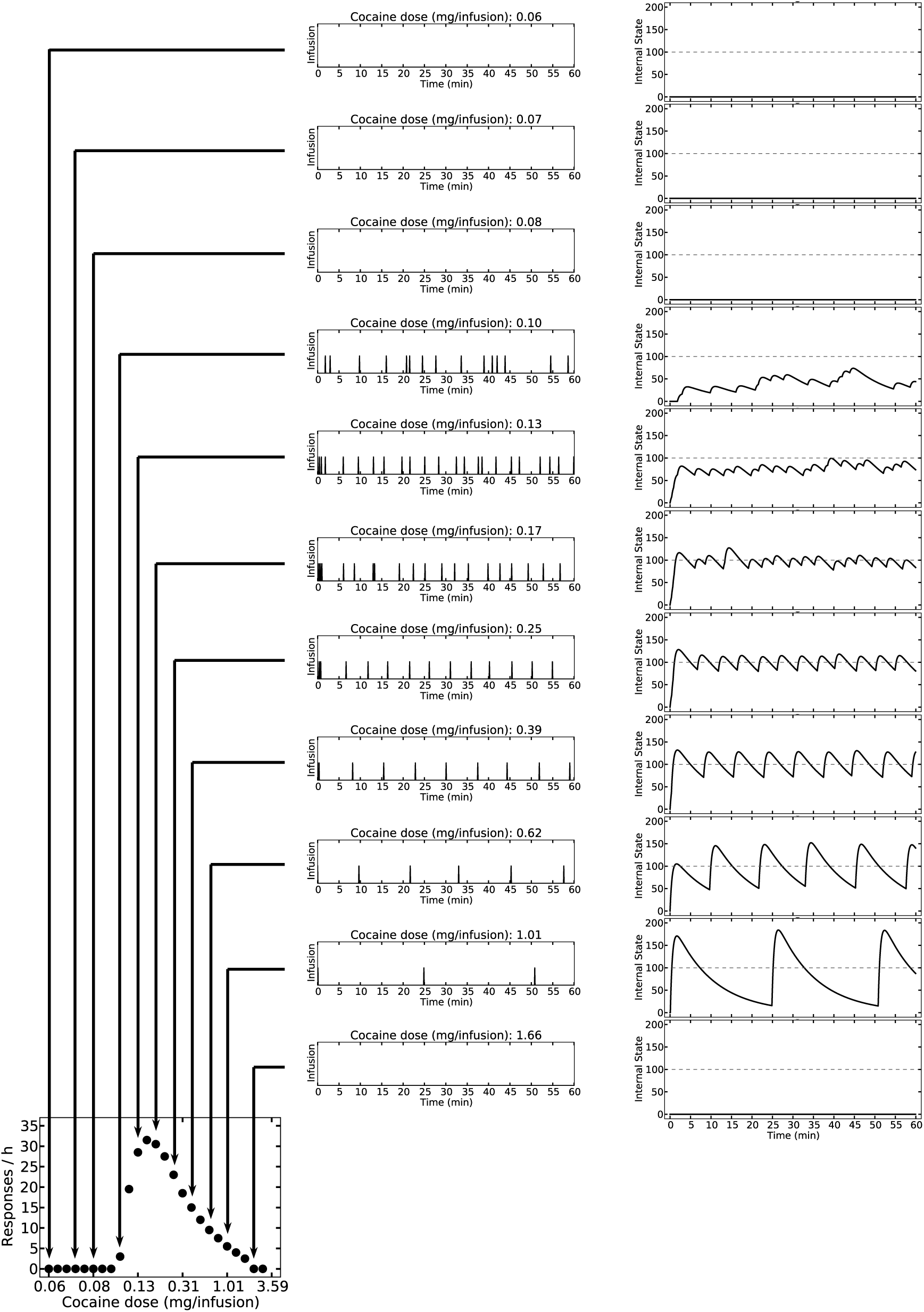
Simulation results showing the underlying mechanism behind the observed dose-response curves. At very low doses, the cost of lever-press outweighs the rewarding value of cocaine. Thus, the rate of self-administration is low. As the dose increases, the rewarding value increases and so does the rate of lever-press. At some point, the unit dose is high enough that its rewarding/motivational value is sufficient for inducing the rate of lever-press necessary for reaching the setpoint. By increasing the dose beyond this critical level, the rate of responding decreases in order to keep the internal state as close as possible to the setpoint. Extremely high doses (the bottom row) result is such huge deviations from the setpoint (possibly life-threatening) that the agent prefers not to take cocaine whatsoever.

**Fig. S17.**
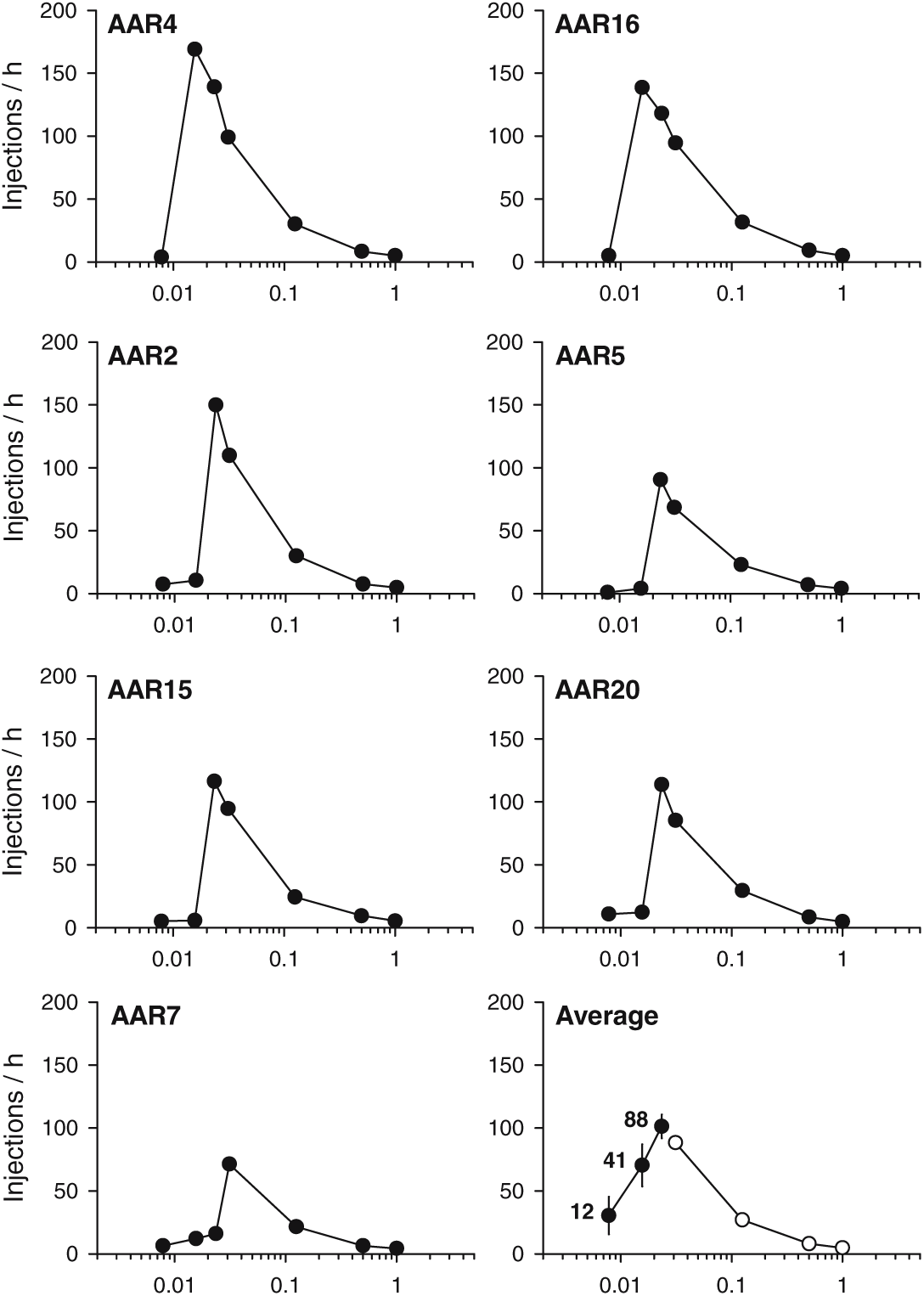
Experimental results (Zittel-Lazarini et al., 2007) showing individual differences in dose-response functions for cocaine self-administration. In all individual curves (for 7 out of n=17 rats), the rate of infusion is negligible at low doses. At a certain dose, the infusion rate peaks and then decreases as a function of the unit dose of cocaine. The critical dose at which infusion rate peaks, as well as the rate of responding at this dose, is different across animals. The bottom-left plots shows the group-average dose–injection function across all individuals (n=17).

**Fig. S18.**
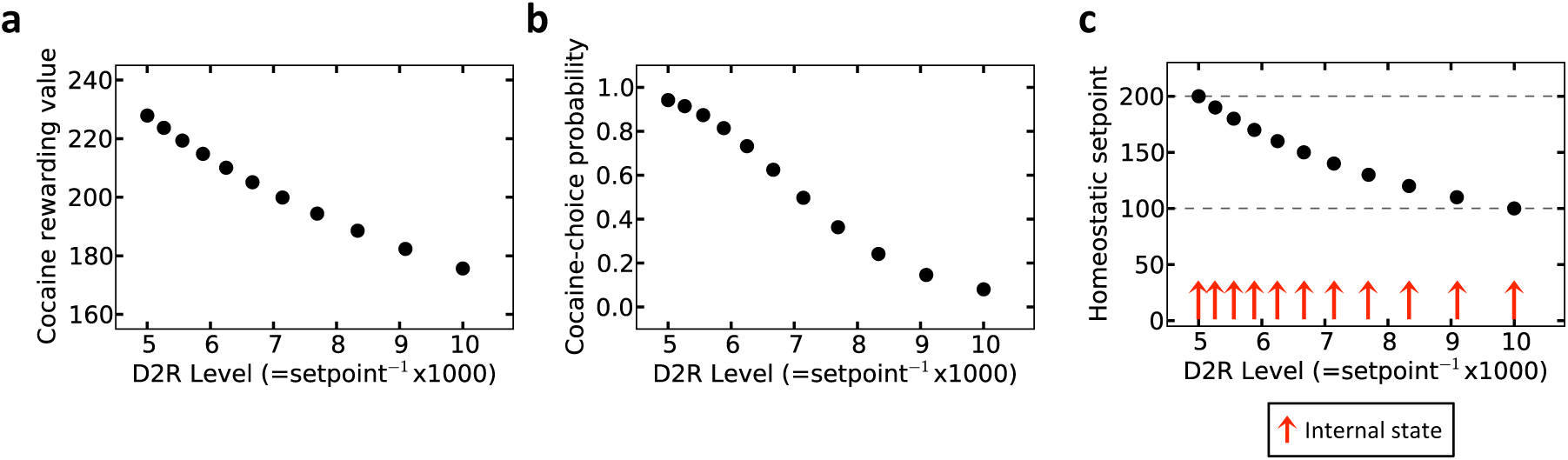
Simulation results replicating experimental data (Volkow et al., 1999) Fig. S19. The rewarding value of a certain unit dose of cocaine decreases as the level of D2 receptor availability increases (a). Thus, when given a choice between drug and food, the probability of choosing the drug outcome is inversely correlated with D2R level (b). Homeostatic setpoint level is assumed to be encoded inversely by D2R availability (plot c; *D2R level* = 1000. *setpoint*^−1^). The highest and lowest levels of the setpoint are 200 and 100, respectively (equivalent to the D2R levels of 5 and 10, respectively). Choosing the drug option increases the level of the internal state (red arrows) and thus, decreases homeostatic deviation. This drive-reduction reward is higher when the initial distance from the setpoint is higher (i.e., in agent with a high setpoint level, or a low D2R level).

**Fig. S19.**
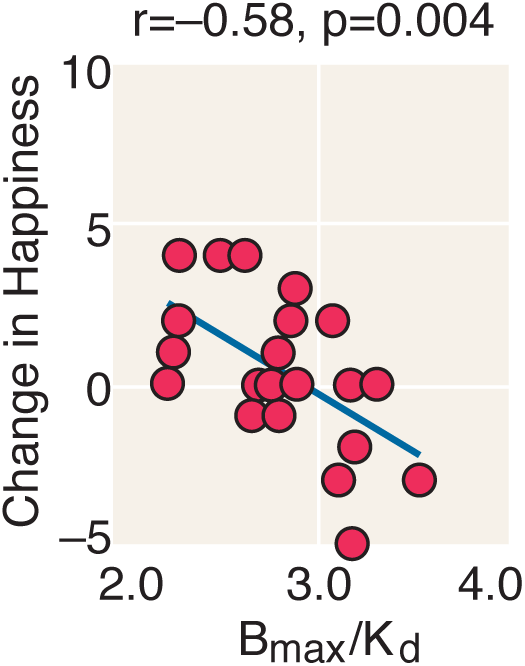
Experimental results (Volkow et al., 1999) showing an inverse correlation between dopamine D2 receptor availability and the reported pleasantness of drug in human subjects with no drug abuse histories.

**Fig. S20.**
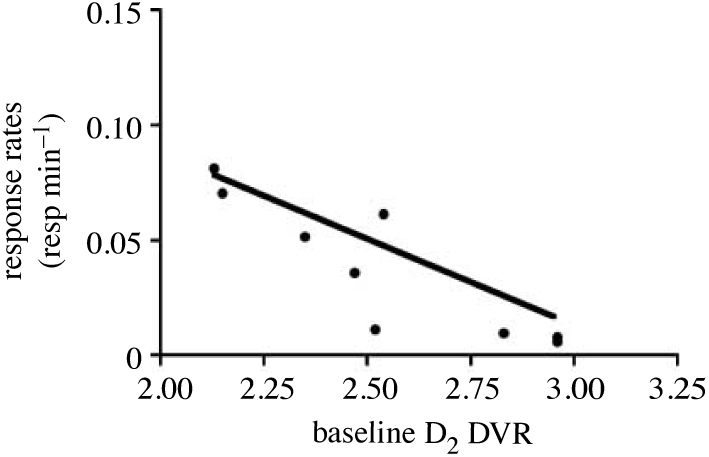
Experimental results (Nader et al., 2006) showing an inverse correlation between dopamine D2 receptor availability in monkeys and the rate of responding for cocaine. Plot reprinted from (Nader et al., 2008).

**Fig. S21.**
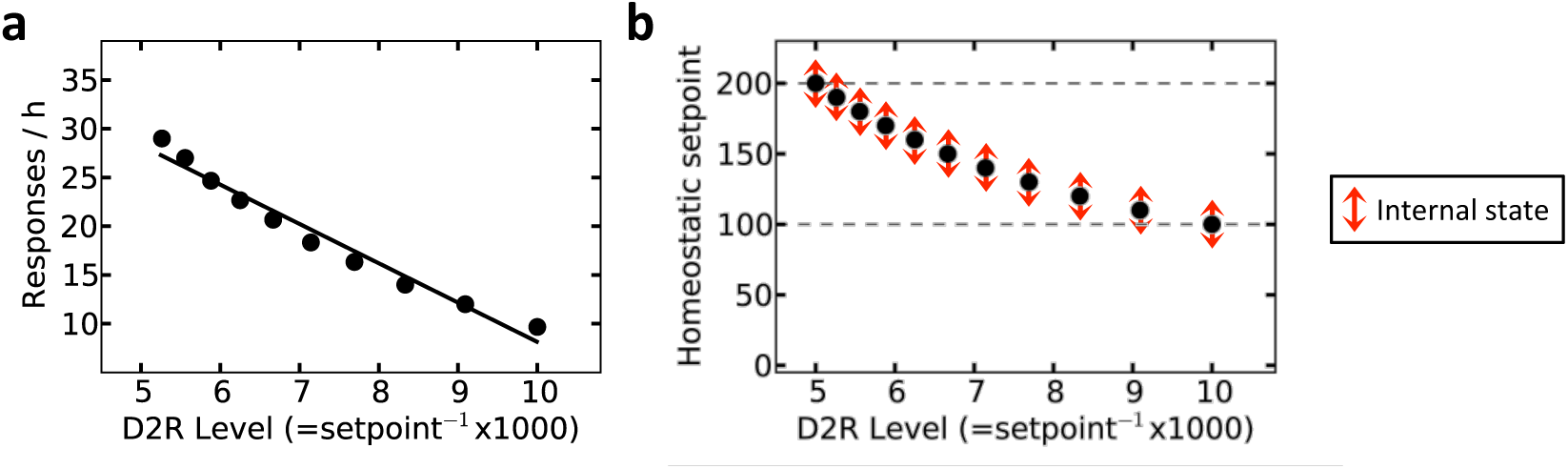
Simulation results replicating experimental data (Nader et al., 2006) (Fig. S20). The rate of cocaine self-administration is inversely correlated with D2R level (a). Homeostatic setpoint level is assumed to be encoded inversely by D2 receptor availability (plot b; *D2R level* = 1000. *setpoint*^−1^). The highest and lowest levels of the setpoint are 200 and 100, respectively (equivalent to the D2R levels of 5 and 10, respectively). For each level of D2R availability, cocaine self-administration results in the internal state fluctuating around the homeostatic setpoint (red arrows).

### Supplementary Tables

**Table. S1.** Free parameters of the model and their values in the simulations.

| Parameter | Value | Explanation |
| --- | --- | --- |
| $\alpha$ | 0.2 | Learning rate of the Q-learning algorithm |
| $\beta$ | 0.25 | Exploration rate in the soft-max rule |
| $\gamma$ | 1 | Discount factor |
| $m$ | 3 | Root of the drive function |
| $n$ | 4 | Power of the drive function |
| $\underline{h^*}$ | 100 | Lower-bound of the setpoint |
| $\overline{h^*}$ | 200 | Upper-bound of the setpoint |
| $h_0^*$ | 100 | Initial setpoint |
| $K$ | 50 | Effect of 0.250mg of cocaine on the internal state |
| $\phi$ | 0.007 | Rate of absorption of cocaine |
| $\omega$ | 0.113 | Rate of elimination of cocaine |
| $\mu$ | 0.0018 | Rate of setpoint up-regulation |
| $\rho$ | 0.00016 | Rate of setpoint recovery |
| $cost$ | 1 | Energy-cost of lever-press |
| $punishment$ | 380 | Punishment associated with drug in Fig. 5 |

